# Engineered Cortical Microcircuits for Investigations of Neuroplasticity

**DOI:** 10.1101/2024.06.08.598052

**Authors:** Nicolai Winter-Hjelm, Pawel Sikorski, Axel Sandvig, Ioanna Sandvig

## Abstract

Recent advances in neural engineering have opened new ways to investigate the impact of topology on neural network function. Leveraging microfluidic technologies, it is possible to establish modular circuit motifs that promote both segregation and integration of information processing in the engineered neural networks, similar to those observed *in vivo*. However, the impact of the underlying topologies on network dynamics and response to pathological perturbation remains largely unresolved. In this work, we demonstrate the utilization of microfluidic platforms with 12 interconnected nodes to structure modular, cortical engineered neural networks. By implementing geometrical constraints inspired by a Tesla valve within the connecting microtunnels, we additionally exert control over the direction of axonal outgrowth between the nodes. Interfacing these platforms with nanoporous microelectrode arrays reveals that the resulting laminar cortical networks exhibit pronounced segregated and integrated functional dynamics across layers, mirroring key elements of the feedforward, hierarchical information processing observed in the neocortex. The multi-nodal configuration also facilitates selective perturbation of individual nodes within the networks. To illustrate this, we induced hypoxia, a key factor in the pathogenesis of various neurological disorders, in well-connected nodes within the networks. Our findings demonstrate that such perturbations induce ablation of information flow across the hypoxic node, while enabling the study of plasticity and information processing adaptations in neighboring nodes and neural communication pathways. In summary, our presented model system recapitulates fundamental attributes of the microcircuit organization of neocortical neural networks, rendering it highly pertinent for preclinical neuroscience research. This model system holds promise for yielding new insights into the development, topological organization, and neuroplasticity mechanisms of the neocortex across the micro- and mesoscale level, in both healthy and pathological conditions.

## Introduction

The underlying microcircuit motifs and architectures of neural networks play a pivotal role in facilitating efficient information processing, transfer, and storage within the brain (1). However, our understanding of how these structural motifs that emerge during brain development contribute to efficient neural computations in physiological conditions, or influence the effect of pathological perturbations, remains incomplete. This knowledge gap is, in part, attributed to the inherent challenges of studying brain network dynamics at the micro- and mesoscale level *in vivo*. Engineered neural networks offer a complementary approach to *in vivo* models by enabling investigation of neuroplasticity in a controlled microenvironment *in vitro*. Engineered neural networks are based on the inherent property of dissociated neurons to self-organize over time into complex computational systems, recapitulating fundamental characteristics of brain networks (2, 3). This process occurs under the influence of various chemically and physically regulated guidance cues from the microenvironment that operate in a spatiotemporal manner in tandem with inherent self-organizing properties of neurons and their spontaneous electrical activity (4–7). The presence of topological cues within the neurons’ microenvironment is highly relevant for the emergence of microcircuit motifs and architectures that mimic the modular organisation of neural circuits of interest, thereby further improving the physiological relevance of the model. Current technologies, such as microfluidics, enable precise manipulation of network topology. Such platforms promote establishment of modular networks, where populations of neurons are segregated into distinct chambers connected by micrometer-sized tunnels accessible only to their neurites (8). Recent studies have shown the importance of modularity in inducing complex dynamics, balancing integrated and segregated activity akin to that seen across brain regions *in vivo* (9–12). By implementing geometrical constraints within the microtunnels, it is furthermore possible to precisely control the direction of axonal outgrowth between the distinct populations of neurons (13–17). This approach, in combination with microelectrode array (MEA) interfaces enabling electrophysiological recordings, can be utilised to recapitulate feedforward microarchitectures observed in many brain areas. In this way, anatomically relevant microcircuits, such as the cortical-hippocampal projection, can be established, thereby providing a robust methodology for studying neural network function and dysfunction in a controlled setting (11, 18).

A better understanding of how the underlying structure of neural networks shapes their dynamics is critical for deciphering physiological behaviours, as well as network responses to pathological perturbations (19). Graph theory has emerged as a prominent mathematical approach for gleaning meaningful insights into neural network connectivity and function (20, 21). Within this framework, several hallmarks of efficient neural network organization have been identified, including small-world properties with high local clustering and short path length between nodes, and a modular, hierarchical network architecture that promotes efficient information flow between distinct parts of the network (21, 22). Furthermore, efficient neural circuits typically feature a small number of highly interconnected hubs, which are critical for integrating information across distinct network regions (23). These topological traits, believed to be genetically encoded in the neurons for instructing neural self-organization, are hypothesized to minimize the metabolic cost of the network while maintaining efficient information flow (24–26). Analyzing and recognizing such traits can provide insights into the computational complexity of neural networks in both *in vivo* and *in vitro* settings (27–30).

Neocortical microcircuits, characterized by their welldefined, layered input/output structure, is a compelling system to engineer using modular, directional microfluidic platforms (31). The neocortex represents one of the most intricate structures in the human brain, being responsible for a broad spectrum of functions, ranging from storage of working memory and predictive coding to various cognitive and sensorimotor tasks (32–36). While exhibiting diverse functions across distinct cortical regions, the entire neocortex is theorized to comprise canonical elementary processing units known as minicolumns (37–40). The precise function of such minicolumns in cortical processing is still under debate, yet they are believed to represent fundamental units of cortical organization. Furthermore, cortical networks exhibit complex spatiotemporal activity patterns during development, even in the absence of external input, both *in vivo* and *in vitro* (2, 3, 13, 41–44). Additionally, cortical circuits demonstrate a remarkable ability to undergo plasticity changes during development and in response to external stimuli or damage (45–47). Thus, the layered structural organization, intricate spontaneous functional dynamics and extensive plasticity of neocortical microcircuits render them compelling motifs for investigation under controlled conditions *in vitro*, offering valuable insights into the structure-function relationships of such circuits.

In this study, we present an advanced microfluidic MEA model for recapitulating the feedforward, hierarchical architecture of the neocortex *in vitro*. Furthermore, we demonstrate the utility of this model system for studying network plasticity in response to selective perturbation of hub nodes that are central for information flow between higher and lower nodes in the structured hierarchy. We evaluate changes in the networks’ information processing by analyzing both spontaneously evoked and stimulation-induced activity and apply connectomics and graph theory to study alterations in information flow within the networks. We thus present an engineered cortical microcircuit model that facilitates the investigation of developmental network dynamics, as well as its behaviour under physiological and pathological conditions.

## Materials and Methods

### Design & Fabrication of Microdevices

The design of the microfluidic MEA was created using Clewin 4 (WieWeb Software, Enschede), as depicted in **Figure S1**. The platform was organized hierarchically into four distinct layers with a feedforward architecture consisting of 12 interconnected chambers, referred to as nodes. Each node had a diameter of 3.5 mm. Two and two nodes were interconnected by 20 microtunnels, each 350 µm long, 10 µm wide and 5 µm high. These channels had geometries inspired by the Tesla valve to induce unidirectional axonal outgrowth between the nodes (13). Additionally, spine structures were integrated on the postsynaptic side to misguide any outgrowing axons trying to enter the tunnels in the unintended direction (17). The microelectrode arrays (MEAs) were engineered to be compatible with a MEA2100 workstation from Multichannel Systems. Electrodes, measuring 2 mm in length and 10 µm in width, were strategically positioned across each channel area, as well as at the entrances and exits of each tunnel area, to monitor all activity propagation between the nodes. Furthermore, two electrodes were placed in each of the nodes in layer 1 and layer 4, with an additional electrode positioned in node 4. All electrodes were functionalized with a thin layer of highly nanoporous platinum to enhance their signal-to-noise ratio. A reference electrode was also positioned in each of the 12 nodes, with all reference electrodes connected to the same channel on the recording system. The fabrication of all microelectrode arrays followed our recently reported protocol (13).

### Coating, Cell Plating and Maintenance

An overview of the experimental timeline is depicted in **Figure 1A**. Prior to coating, all interfaces underwent sterilization in UV light for a minimum of 1 h. Subsequently, the samples were soaked in DMEM, low glucose (Gibco™, 11885084) for at least 48 h to remove any potentially toxic, uncured PDMS left in the microfluidic chips. The interfaces were then coated with 0.1 mg*/*mL Poly-L-Ornithine solution (PLO) (Sigma-Aldrich, A-004-C) overnight in a fridge at 4 ^°^C. The following day, all chambers were washed three times with Milli-Q (MQ) water to remove unattached PLO, before coating the surfaces with a laminin solution consisting of 16 µg*/*mL natural mouse laminin (Gibco™, 23017015) diluted in phosphate-buffered saline (PBS, Sigma-Aldrich, D8537) at 37 ^°^C, 5 % CO_2_ for 2 h. To ensure flow of the coating solution through the microtunnels, a hydrostatic pressure gradient was established between the chambers by filling them with varying amounts of the solution.

**Figure 1.**
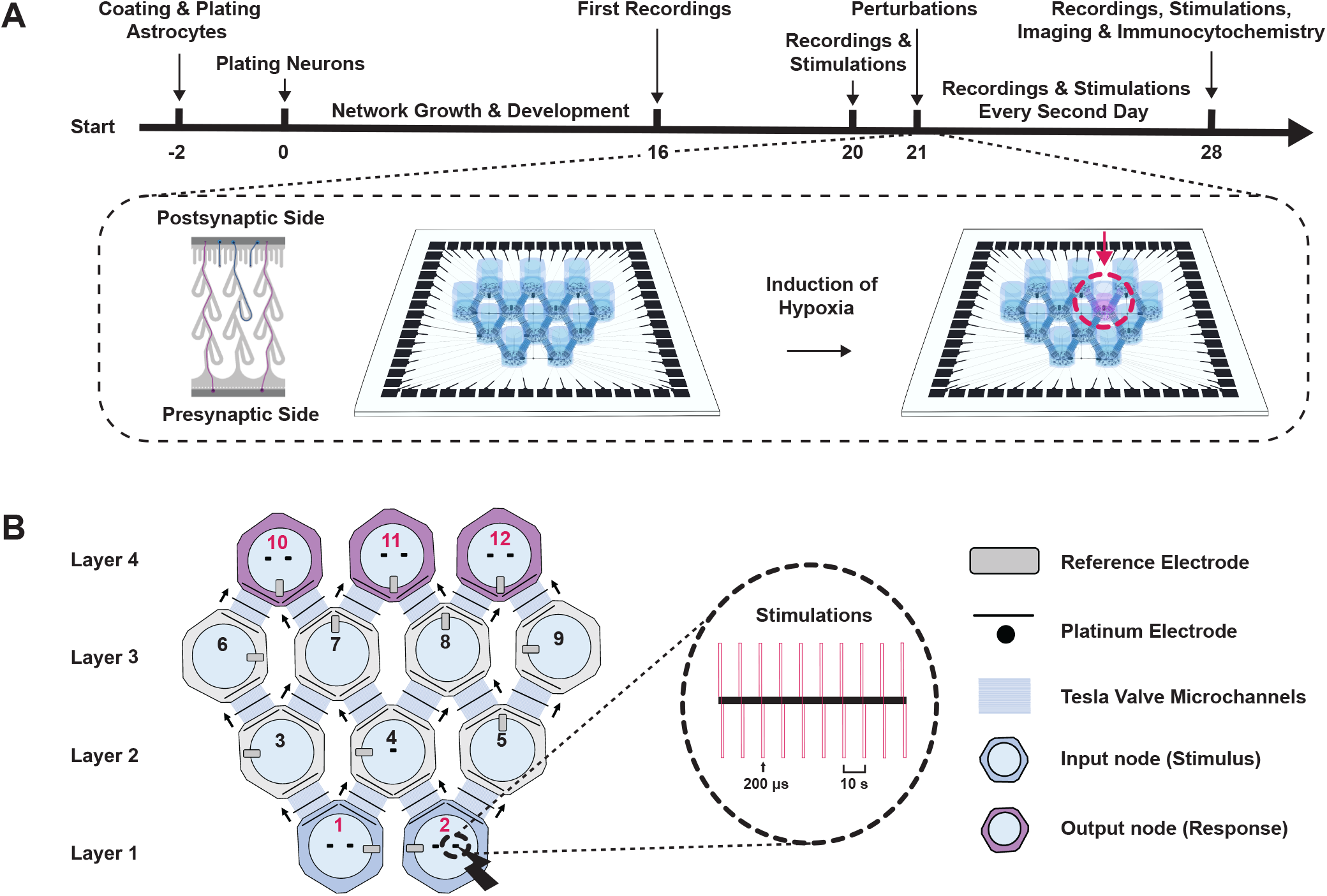
Experimental setup. **(A.)** Experimental timeline for cell experiments. An astrocytic feeder layer was plated two days prior to plating of neurons. Baseline recordings were conducted at 16 and 20 days *in vitro* (DIV). At 21 DIV, localized hypoxia was induced in one of the central nodes in layer 3 using CoCl_2_. Electrophysiological recordings were subsequently conducted every second day until 28 DIV. **(B.)** Schematic illustrating the organization of the 12 nodes, arranged into four distinct layers containing 2, 3, 4, and 3 nodes, respectively. Stimulations were administered following the recordings of spontaneously evoked activity from 20 to 28 DIV. Each stimulation session consisted of 10 consecutive spikes with a 10 s interspike interval and was applied to all input and output nodes (nodes 1, 2, 10, 11 & 12) in increasing order based on their node numbers. The illustrations are not drawn to scale.

An astrocytic feeder layer was plated two days prior to plating of neurons. Astrocyte medium consisted of DMEM, low glucose supplemented with 15 % Fetal Bovine Serum (Sigma-Aldrich, F9665) and 2 % Penicillin-Streptomycin (Sigma-Aldrich, P4333). Rat astrocytes (Gibco™, N7745100) were plated at a density of 75 cells*/*mm^2^, equivalent to 750 cells per culturing chamber. The cells were allowed to expand for two days before the plating of neurons. The astrocyte medium was subsequently replaced by neuronal medium consisting of Neurobasal Plus Medium (Gibco™, A3582801) supplemented with 2 % B27 Plus (Gibco™, A358201), 1 % GlutaMax (Gibco™, 35050038) and 2 % Penicillin-Streptomycin (Sigma-Aldrich, P4333). 0.1 % Rock Inhibitor (Y-27632 dihydrochloride, Y0503, Sigma-Aldrich) was included in the medium during plating to enhance cell viability. Sprague Dawley rat cortical neurons (Gibco, A36511) were plated at a density of 750 cells*/*mm^2^, i.e., 7500 cells per chamber. Following this, half the cell medium was replaced with fresh neuronal medium at 4 h and 24 h after plating. Subsequently, the medium was replaced every second day throughout the experimental period. All electrophysiology experiments were conducted using cells from the same batches and cell vials.

### Immunocytochemistry

For immunocytochemistry, cells were plated in microfluidic platforms bonded to glass coverslips (VWR International, 24×24 mm No. 1 Menzel-Gläser). Prior to fixation, cells were washed with PBS to remove debris. Fixation was performed for 15 min at room temperature using glyoxal solution comprising 20 % absolute ethanol (Kemetyl, 100 %), 8.0 % Glyoxal solution (Sigma-Aldrich, 128465), and 1 % acetic acid (Sigma-Aldrich, 1.00063) in MQ-water (48). Following fixation, the cells underwent three consecutive PBS washes, each lasting 15 min. Subsequently, cells were permeabilized with 0.5 % Triton-X (Sigma-Aldrich, 1086431000) diluted in PBS, followed by two additional PBS washes to remove excess Triton-X. The cells were then blocked with a solution consisting of 5 % goat serum (Abcam, ab7481) diluted in PBS and incubated at room temperature on a shaking table at 30 rpm for 1 h. Primary antibody solutions, prepared in PBS with 5 % goat serum as well as antibodies at concentrations listed in **Table 1**, was added to the cells. The cultures were then incubated overnight on a shaker table at 30 rpm at 4 ^°^C. The following day, cells were rinsed three times with PBS for 15 min each, followed by incubation with 0.2 % secondary antibodies diluted in PBS and 5 % goat serum at room temperature for 3 h on a shaker table at 30 rpm. Prior to application, the secondary antibody solution was centrifuged at 6000 rpm for at least 15 min to remove precipitates. Subsequently, 0.1 % Hoechst (Abcam, ab228550) diluted in PBS was added to the cultures, and the cultures were incubated for an additional 30 min on the shaker table. Before imaging, all cultures underwent three PBS washes followed by two rinses in MQ water.

**Table 1.** Antibodies and concentrations used for Immunocytochemistry.

| Marker | Catalogue Number | Concentration |
| --- | --- | --- |
| GFAP | Ab7260 | 1/1000 |
| NeuN | Ab279295 | 1/500 |
| Neurofilament Heavy | Ab4680 | 1/5000 |
| PSD95 | Ab13552 | 1/250 |
| Synaptophysin | Ab32127 | 1/500 |

*All antibodies were purchased from Abcam.*

### Viral Transductions for Structural Characterization

To induce ubiquitous expression of GFP and mCherry within distinct nodes, viral transductions were performed using an AAV 2/1 serotype construct containing either pAAV-CMV-beta Globin intron-EGFP-WPRE-PolyA or pAAV-CMV-beta Globin intron-mCherry-WPRE-PolyA plasmids driven by a CMV promoter. The viral vectors were produced in-house by Rajeevkumar Raveendran Nair at the Viral Vector Core Facility, NTNU. Transductions were conducted at 9 DIV (n = 4). Initially, 3/4 of the neuronal media in the nodes located in layers 1 and 3 of the 12-nodal microfluidic chips were aspirated, and viruses for expression of GFP were introduced at a concentration of 5*e*^2^ viruses/cell. After a 3 h incubation period, the transduced nodes were replenished with fresh media. Subsequently, 3/4 of the media in the nodes situated in layers 2 and 4 were removed, and viruses for expression of mCherry were applied at a concentration of 5*e*^2^ viruses/cell. Following another 3 h of incubation, these node were also replenished with fresh media. Network imaging was conducted at 21 DIV (prior to induction of hypoxia) and at 28 DIV.

### Chemical Perturbation to Induce Hypoxia

Cobolt chloride (CoCl_2_) (Merck, 15862) was used to induce hypoxia in either node 7 or 8 at 21 DIV (49). The node demonstrating the greatest functional connectivity among the electrodes within the incoming and outgoing microtunnels of the node at 21 DIV was selected for perturbation. Initially, 3/4 of the media in the targeted node were removed, followed by the addition of fresh cell media containing 1000 µM CoCl_2_. To keep the impact of the perturbation localized to a single node, the chamber was filled only up to half its volume, thereby establishing a hydrostatic pressure gradient between the targeted and neighboring nodes. This yielded an overall concentration of 500 µM CoCl_2_ in the targeted node.

### Calcium Imaging

To conduct calcium imaging, the cells in 6 microfluidic chips bonded to glass coverslips (VWR International, 24×24 mm No. 1 Menzel-Gläser) were transduced with in-house prepared viral vectors with a AAV8 serotype construct containing pAAV.CAG.GCaMP6s.WPRE.SV40 plasmids (Addgene, 100844). The cells were transduced at 17 DIV by replacing the cell medium with fresh neuronal medium containing a viral load of 5*e*^2^ viruses/cell. Imaging was conducted between 20 - 28 DIV.

### Imaging

Fluorescence microscopy and calcium imaging were performed using either an EVOS M5000 microscope (Invitrogen) or an EVOS7000 microscope with an onstage incubator (Invitrogen). DAPI (AMEP4650), CY5 (AMEP4656), GFP (AMEP4651) and TxRed (AMEP4655) LED light cubes and Olympus UPLSAP0 4X/0.16 NA and 20x/0.75 NA objectives were utilized. Post-processing of images was conducted in ImageJ/Fiji or Adobe Photoshop 2020. For the calcium imaging, the microfluidic chips were covered by a custom-designed 3D-printed cap (UltiMaker Cura 3D printer) covered with a gas permeable membrane (MEA Membrane Cover, ALA scientific instruments). Recordings were conducted for 2 minutes at 3 frames/s.

### Electrophysiological Recordings

Electrophysiological recordings were performed using a MEA2100 workstation (Multichannel Systems) with the sampling rate set to 25000 Hz. A constant temperature of 37 ^°^C was maintained using a temperature controller (TC01, Multichannel Systems). Sterility of the cultures during recordings was ensured by employing a 3D-printed plastic cap with a gas permeable membrane (MEA Membrane Cover, ALA scientific instruments). Prior to recordings, the neural networks were allowed to equilibrate for 5 min on the recording stage, followed by a 15 min recording period. All recordings were conducted 24 h after media changes.

### Electrical Stimulations

Stimulations were applied subsequent to the recordings of spontaneous network activity on 20, 22, 24, 26 and 28 DIV. The stimulation protocol applied a series of 10 consecutive pulses at *±* 800 mV amplitude (positive phase first), each lasting 200 µs, with an interspike interval of 10 s (**Figure 1B**). Nodes 1, 2, 10, 11 and 12 in layers 1 and 4 of the 12-nodal networks were selected for stimulation to verify the establishment of functional feedforward microcircuits. Stimulations were applied to the nodes in increasing order. Each spike train was delivered to the most active electrode positioned at the center of the targeted node, determined by the firing rate of the electrodes.

### Data Analysis Calcium Imaging

For analysis of the calcium imaging recordings, the open-source software CAL- IMA was utilized (50, 51). Downscaling was set to 2x to shorten execution time prior to analysis. The standard deviation was set to 2.5, 3.5 and 0.003 for the three Gaussian filters employed by the software to detect regions of interest (ROI), respectively. Spike detection was based on the mean values (ΔF/F0) per ROI, and parameters were set to 10 s windows, a Z-score of 3.0 and an *m* value of 0.6. Matlab R2021b was used for further analysis. The *SpikeRasterPlot* function developed by Kraus (52) was adapted and used for creating raster plots.

### Data Analysis Electrophysiology

Data analysis was performed using Matlab R2021b, with graphs plotted using the linspecer function, based on the colorBrewer palette (53, 54). To preprocess the data, a 4th-order Butterworth bandpass filter was applied to remove frequencies below 300 Hz and above 3000 Hz. Additionally, noise from the power supply mains at 50 Hz was eliminated using a notch filter. Zerophase digital filtering was employed to avoid group delay in the output signal. Spike detection was carried out using the Precise Timing Spike Detection (PTSD) algorithm developed by Maccione *et al*. (55). The data was thresholded at 8 times the standard deviation of the noise, with maximum peak duration and refractory time set to 1 ms and 1.6 ms, respectively. The *SpikeRasterPlot* function developed by Kraus (52) was adapted to create raster plots.

After binning the data into 50 ms time bins, functional connectivity was analyzed using Pearson correlation. To visualize key network features, graphs were plotted with electrodes as nodes and Pearson correlation as edges. Edges with connectivity less than 0.05 were removed to highlight only the strongest connections. Node color was used to represent firing rate (Hz), while node size corresponded to PageRank centrality. Community detection was performed using the Louvain algorithm, which delineated nodes into distinct communities based on the strength of their interconnections (56). Node edge colors in the graphs indicated community membership. Graph theoretical measures represented in the graphs were calculated using the Brain Connectivity Toolbox developed by Rubinov & Sporns (57).

To calculate the path length between nodes in layers 1 and 4 of the networks, the inverse of the summed correlation between nodes was utilized. Specifically, the correlation for distinct paths was determined by summing the correlations between nodes/tunnels along the path. For example, for path 1 going from node 1 to 10, a sum of the correlation between the electrodes in node 1 and tunnel 1, tunnel 1 and tunnel 2, tunnel 2 and tunnel 3 and tunnel 3 and node 10 was used. If one or more correlation values were zero, the path length was considered infinite and the data point not included in the graph. The total path length between two nodes was calculated by summing the path lengths of all available paths connecting the nodes.

To remove stimulation artifacts, the stimulation data was processed using the SALPA filter developed by Wagenaar *et al*. (58). Additionally, 15 ms of filtered data was blanked following each stimulation time point. The data was then binned into 30 ms time intervals, and peristimulus time histograms (PSTHs) of the average response of stimulations in each tunnel were plotted. Tunnels were considered active if they exhibited an integrated average response of at least 10 spikes within the first 300 ms following the stimulations, which served as the threshold for evaluating stimulation-evoked propagation of activity across the layers. All PSTHs were additionally manually inspected to identify false negatives and false positives. This decision was based on the presence of clearly distinguishable peaks and whether these peaks followed the correct order according to the sequence of the channels between the nodes.

## Results

### The 12-Nodal Microfluidic Devices Facilitate Establishment of Feedforward, Hierarchical Cortical Microcircuits

To assess the efficacy of the 12-nodal microfluidic platform in fostering the development of mature feedforward, hierarchical neural microcircuits, structural characterization was conducted using immunocytochemistry, optical, and fluorescence microscopy. By 16 DIV, neurons had organized into densely connected networks within the individual nodes (**Figure 2A**) and established long-range connections across the microtunnels interconnecting the nodes (**Figure 2B**). Geometric features inspired by the Tesla valve design were integrated into the microtunnels to promote unidirectional axonal outgrowth between the nodes, complemented by saw-tooth structures on the postsynaptic side to prevent axons from finding the inlets (13). The application of viral tools facilitated the expression of different fluorescent probes in distinct nodes of the multi-layer cortical networks. This confirmed the effectiveness of the Tesla valves in redirecting axons from the postsynaptic node back to their node of origin while guiding axons from the presynaptic node to the opposite node (**Figure 2C**). Furthermore, immunocytochemistry validated the structural maturation of the networks within the nodes by 21 DIV, with the markers Neural Nuclear Protein (NeuN) and Neurofilament Heavy (NFH) (**Figure 2D**) (59, 60). Additionally, GFAP staining was employed to identify the astrocytic feeder layer (61). Colocalization of the pre- and postsynaptic markers synaptophysin and PSD95 confirmed the presence of mature synaptic connections in the networks (**Figure 2D**) (62, 63). Overall, this characterization confirmed structural and functional maturity, with networks exhibiting a feedforward, hierarchical architecture across the distinct layers and nodes.

**Figure 2.**
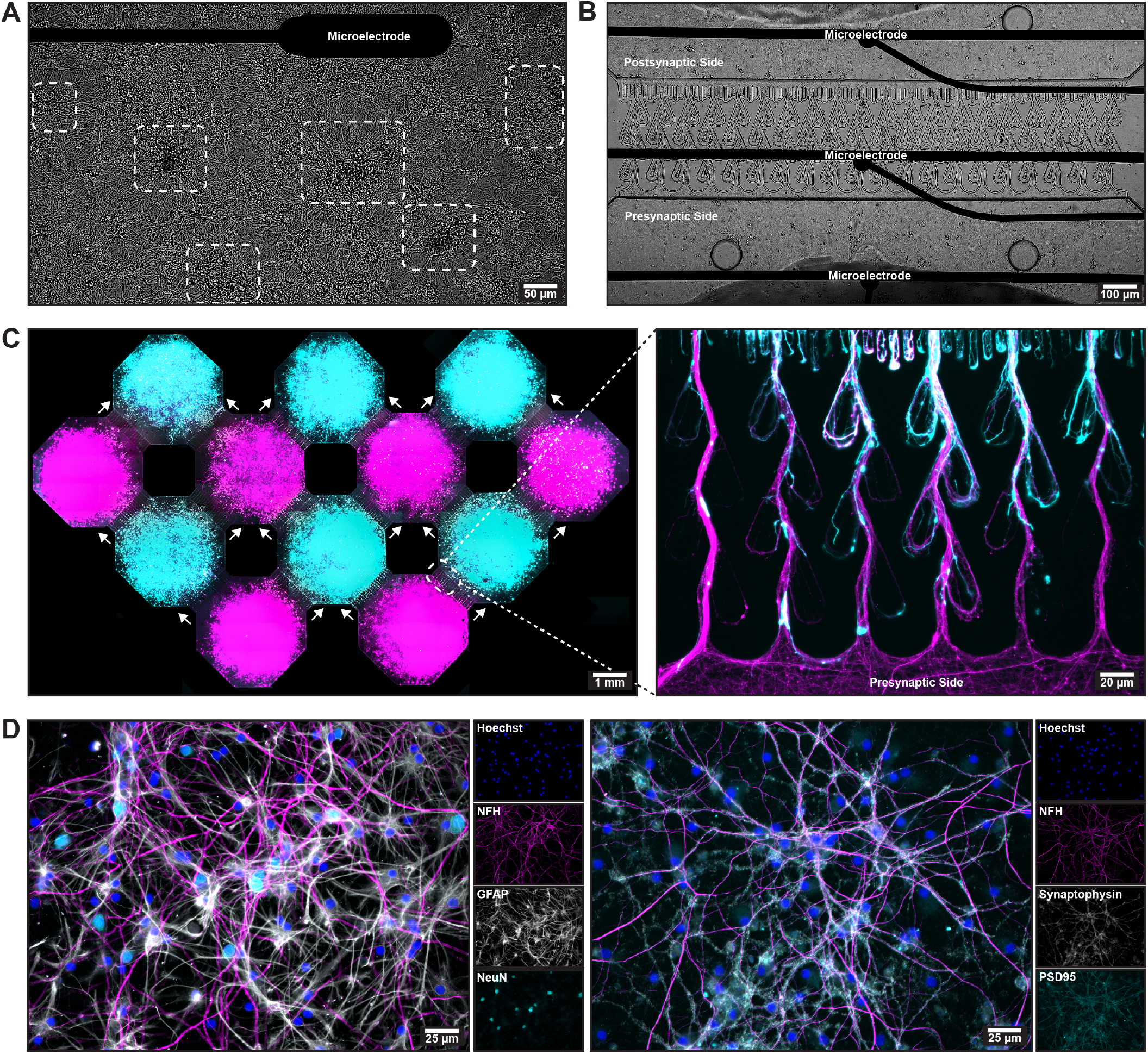
Establishment of mature feedforward, hierarchical neural microcircuits within the 12-nodal platforms. **(A.)** Micrograph showing a neural network growing on top of a microelectrode array (MEA) at 21 days *in vitro* (DIV). The white boxes outline aggregated clusters of neurons. **(B.)** Micrograph illustrating the outgrowth of axons through the Tesla valve microtunnels at 21 DIV. Three electrodes can be seen spanning the width of the tunnels to capture neural activity propagating across the two displayed nodes. **(C.)** Micrographs of a 12-nodal network with neurons expressing either GFP (layer 1 and 3, magenta) or mCherry (layer 2 and 4, cyan) at 28 DIV. The images show how the Tesla valve microfluidic tunnels promote feedforward axonal outgrowth, by rerouting axons growing in the unintended direction back to their node of origin. **(D.)** Immunocytochemistry micrograph showing that the neural networks reached a mature state by 27 DIV, indicated by the presence of the mature structural and nuclei markers Neurofilament Heavy (NFH) and NeuN, respectively. **(E.)** Immunocytochemistry micrograph showing the colocalization of the pre- and postsynaptic markers synaptophysin and PSD95, indicating the establishment of mature synapses.

At 16 DIV, electrophysiological recordings unveiled a complex functional interplay of segregated and integrated activity across all network layers and nodes (**Figure 3A**). Networkwide bursts were observed propagating from the bottom layer (layer 1) to the uppermost layer (layer 4) of nodes across the three layers of microtunnels (**Figures 3B & 3C**). Additionally, externally applied electrical stimulations to the nodes in layer 1 of the networks induced activity that propagated across all three layers of microtunnels in more than 50 % of the available pathways. In contrast, stimulations to the nodes in layer 4 did not evoke detectable responses beyond the first layer of microtunnels in over 99 % of available pathways (**Figures 3D - 3F**). 60 % of stimulation-induced activity in layer 4 was detected in the first layer of microtunnels, i.e., between layer 3 and 4. However, it is worth noting that the electrodes detecting this activity were positioned halfway through the tunnels, and is not representative of activity propagating across the full length of the tunnels. Nevertheless, these findings illustrate a defined hierarchical structural and functional organization within the networks, with potent propagation of activity from layer 1 to layer 4, but limited propagation of activity in the opposite direction from layer 4 to the lower layers.

**Figure 3.**
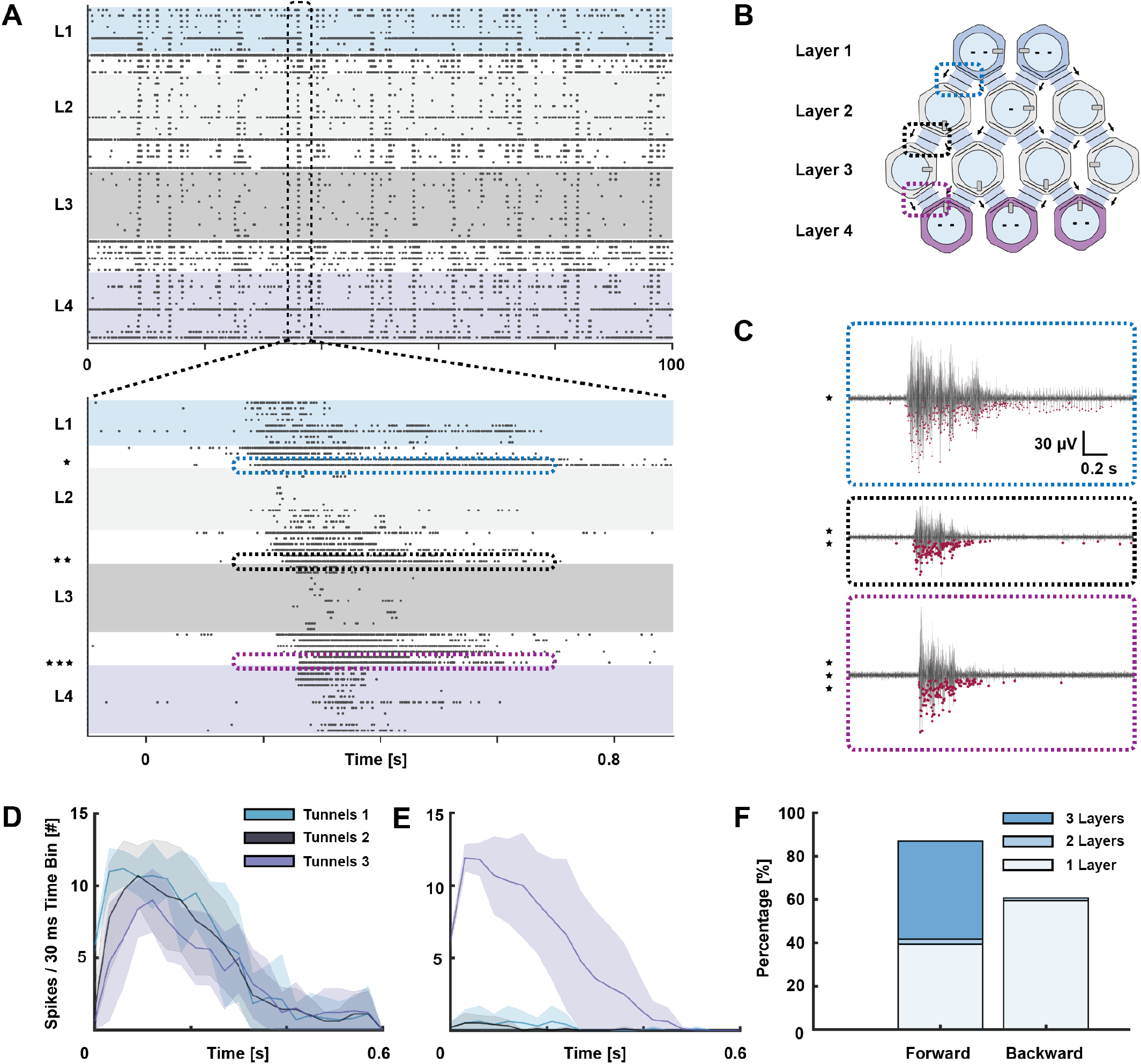
Electrophysiological characterization of information flow and dynamics in the laminar cortical microcircuits. **(A.)** Raster plots showing the complex functional dynamics emerging in a 12-nodal cortical network at 16 days *in vitro* (DIV). A balance between integrated and segregated activity can be seen. A network burst propagating across large parts of the network is outlined with a black box. Furthermore, the propagation of a single network burst from layer 1 to layer 2 (blue box), layer 2 to layer 3 (black box) and layer 3 to layer 4 (purple box) is highlighted. **(B.)** Schematic showing the organization of the 12 nodes into 4 distinct layers, highlighting the positions were the bursts in figure A and C were recorded from. **(C.)** Voltage traces showing the propagation of a single burst detected in the tunnels connecting layer 1 and 2, 2 and 3 and 3 and 4, respectively. **(D.)** - **(E.)** Peristimulus time histograms displaying the average response of 10 consecutive stimulations at each of the three tunnels connecting layer 1 to layer 4 in the hierarchy, when stimulating nodes in layer a and 4, respectively. Stimulations in one of the nodes of layer 1 induced clear responses in the consecutive levels, while no response was observed beyond the first tunnels upon stimulation of the nodes in level 4. **(F.)** Histogram showing percentage of stimulation induced activity spanning 1, 2 or 3 layers when stimulating nodes in layer 1 (forward) or 4 (backward), respectively. While more than 50 % of available pathways exhibited activity propagating across all three layers of microtunnels in the feedforward direction, stimulations to the nodes in layer 4 did not evoke detectable responses beyond the first layer of microtunnels in more than 99 % of available pathways in the backward direction.

In addition to MEA-based electrophysiology, calcium imaging was utilized to examine activity within distinct nodes in greater spatial detail (**Figure 4A**). This technique was employed to investigate whether the multi-nodal design, with multiple inputs to each node, influenced the synchronization of activity within the individual nodes. The calcium imaging revealed high degrees of intranodal synchrony across all layers, including the top layer receiving inputs along multiple pathways (**Figures 4B & 4C**). This finding not only demonstrated the efficacy of the design in promoting coordinated neural activity but also highlighted the potential for combining electrophysiology and calcium imaging on these platforms to study structure-function dynamics with high spatiotemporal precision. Additionally, the observed synchrony both across and within the nodes underscored the functional maturation of the networks, setting a reliable baseline for the perturbations performed at 21 DIV.

**Figure 4.**
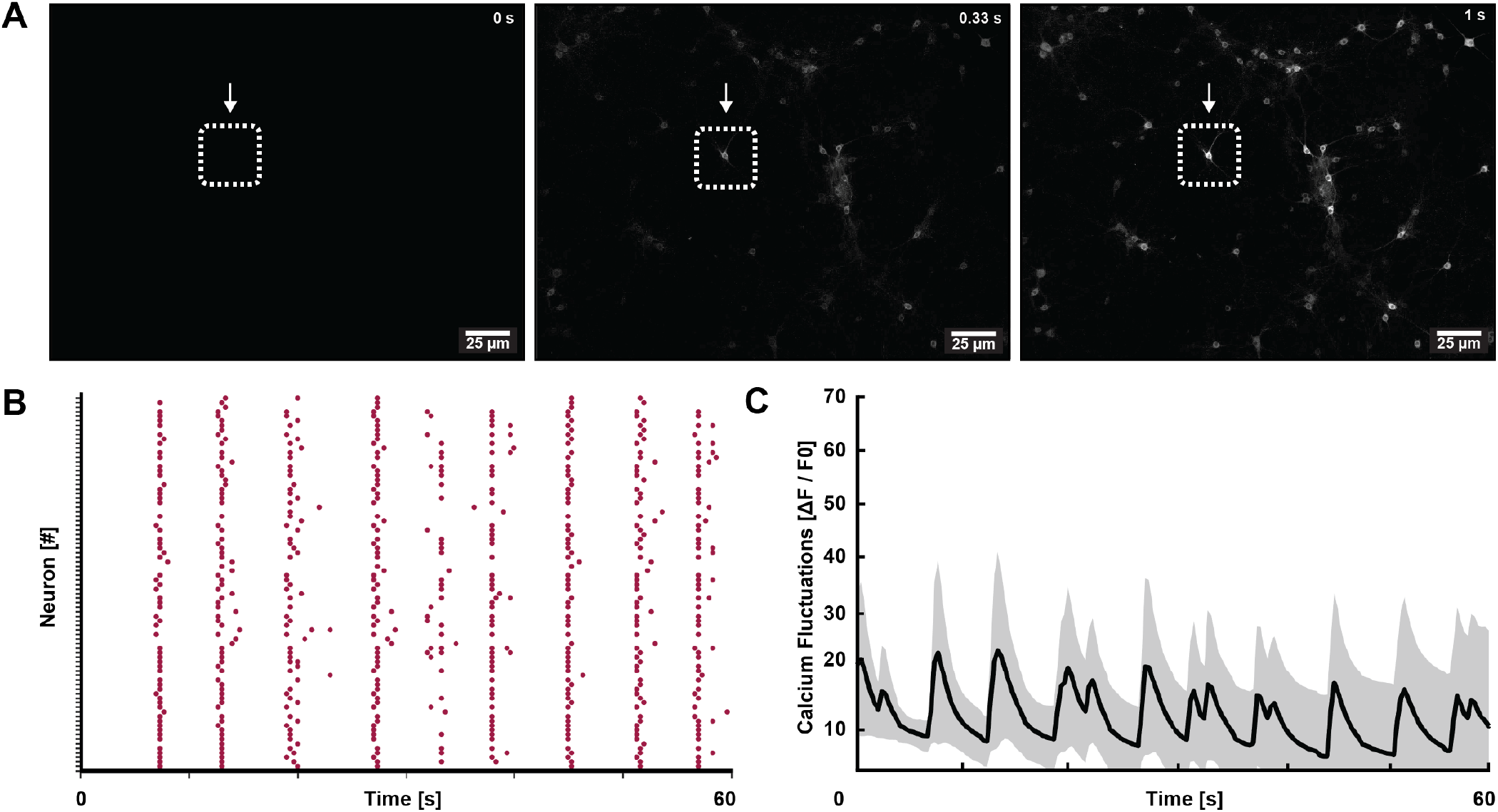
Evaluation of functional intranodal dynamics using calcium imaging. **(A.)** Images captured over a 1 s period showing the progressive increase in fluorescence intensity coinciding with the onset of a network burst. The white boxes highlight a single neuron exhibiting particularly strong calcium fluctuations. **(B.)** Raster plot illustrating the highly synchronized activity of 83 neurons captured within a single field of view during recordings of spontaneously evoked calcium fluctuations in node 11 of a 12- nodal cortical microcircuit at 22 days *in vitro* (DIV). A comparable degree of synchronicity was consistently observed across other nodes in the networks, underscoring their functional maturation. **(C.)** Averaged calcium fluctuations from a single field of view during a 1 min recording at 22 DIV, demonstrating the slow, dynamic temporal calcium fluctuations occurring within the nodes.

### Localized Perturbations Induce Node Ablation and Alters The Network Connectome

To demonstrate the applicability of the model system in studying neuroplasticity in response to localized perturbations, chemical hypoxia was induced in a central node of layer 3 at 21 DIV. Neural networks rely critically on a continuous supply of oxygen for healthy functioning, and hypoxia has been implicated in the pathogenesis of several neurodegenerative diseases (29, 64, 65). In the days following the perturbation, all networks exhibited rapid and extensive fragmentation within the targeted node (**Figures 5A & 5B**). This fragmentation was particularly noticeable towards the center of the targeted nodes, but could also be observed extending towards the microtunnels (**Figure 5C**). Moreover, axons entering the targeted node were observed to retract from the tunnels over time (**Figure 5D**). However, the perturbation did not significantly impact the viability of the neurons within the neighboring nodes (**Figure 5E**). Analysis of electrophysiological activity detected by electrodes in the microtunnels leading to the targeted nodes revealed a rapid functional disconnection of the nodes from the remaining parts of the networks (**Figures 5F & 5G**). These results clearly demonstrate that induction of hypoxia had a detrimental impact on the structural and functional connectivity of the targeted nodes within the first 24 h following perturbation, as intended.

**Figure 5.**
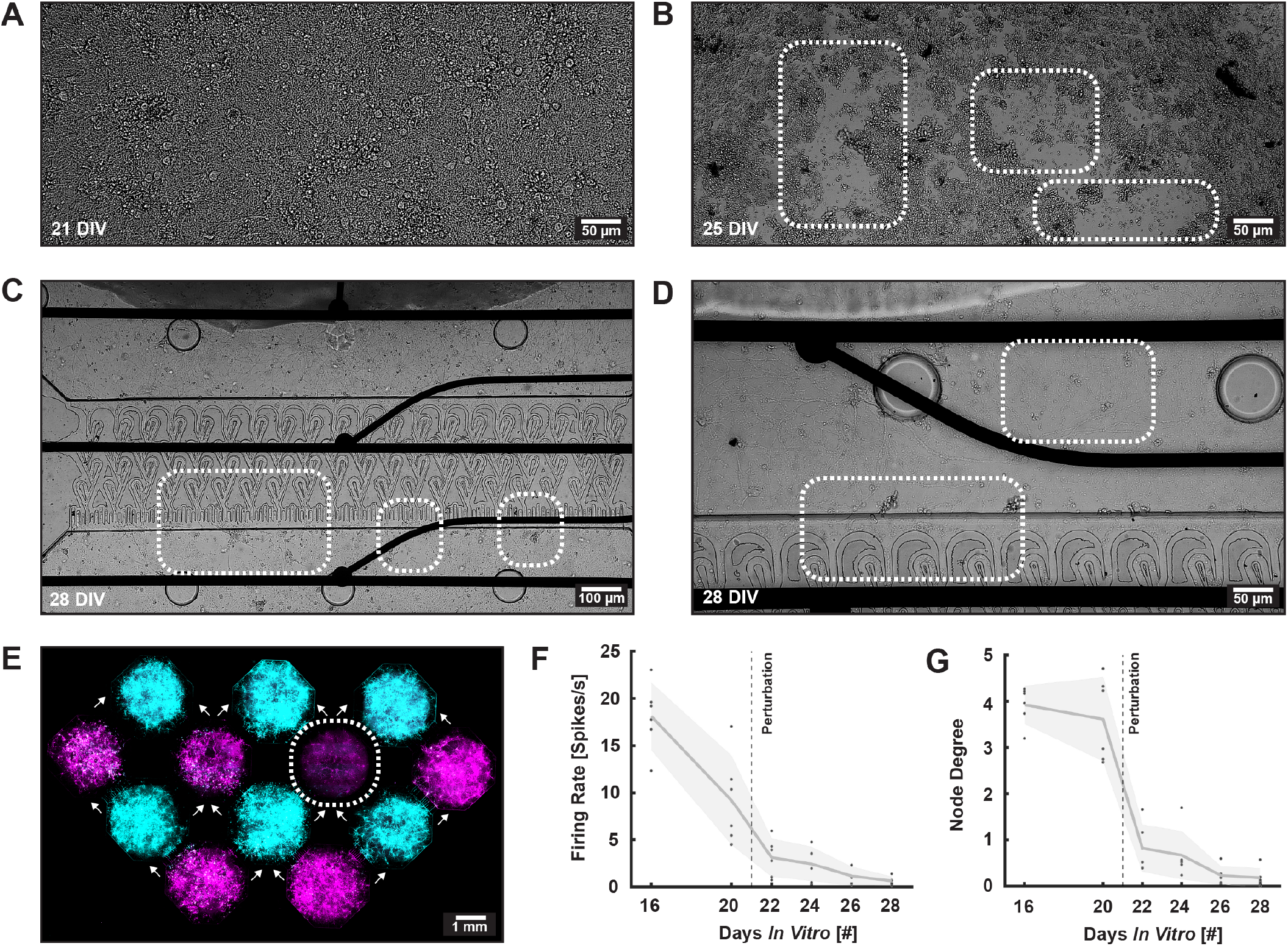
Induction of Localized Perturbation to a hub node. **(A.)** Micrograph depicting a healthy neural network prior to perturbation at 21 days *in vitro* (DIV). **(B.)** Micrograph illustrating the fragmentation of the same neural network 4 days after perturbation. **(C.)** Micrograph displaying the microtunnels connecting a healthy presynaptic node and a perturbed postsynaptic node 7 days after perturbation. The white boxes highlight the fragmentation occuring in the perturbed node. **(D.)** Micrograph showing the presynaptic side of microtunnels leading to a perturbed node, depicting the retraction of neurites growing towards the perturbed node. **(E.)** Micrograph of a 12-nodal network with neurons expressing either GFP (layer 1 and 3, magenta) or mCherry (layer 2 and 4, cyan). The image demonstrates the clear difference in fluorescence intensity within the perturbed node (white box) compared to the surrounding, unperturbed nodes. **(F.)** Firing rate detected by electrodes within the microtunnels connected to the perturbed node, exhibiting a steep decline following perturbation at 21 DIV. **(G.)** Correlation between the activity detected by the electrodes in the microtunnels directly adjacent the perturbed node, indicating the extent to which activity entering the node elicits an outgoing response. A steep decline can be seen following perturbations at 21 DIV.

To investigate dynamic changes in the functional connectome of the networks before and after perturbation, graph theoretical analysis was applied. Initially, at 16 DIV, robust interconnectivity was evident throughout the networks (**Figure 6A**). By 20 DIV, a subtle decrease in functional connectivity was observed, potentially reflecting network pruning and refinement as the networks matured (**Figure 6B**). Based on the recordings at 20 DIV, Pearson correlation was used to identify the most central node in layer 3 for functional integration between layer 1 and layer 4 of each network (outlined with a pink arrow in **Figure 6B**). This node was subsequently selected for perturbation at 21 DIV. By 24 DIV, 3 days following the perturbation, functional connectivity across the targeted node was markedly disrupted, while functional connections in unperturbed regions of the networks appeared strengthened (**Figure 6C**). By 28 DIV, further refinement and alterations in the connectome were apparent, as evidenced by the differential delineation of the networks into distinct communities using the Louvain algorithm (**Figure 6D**). In summary, the application of graph theory revealed dynamic alterations in the functional connectome of the 12-nodal neural networks following localized perturbation, highlighting the intricate adaptive responses and network reorganization processes in response to perturbation.

**Figure 6.**
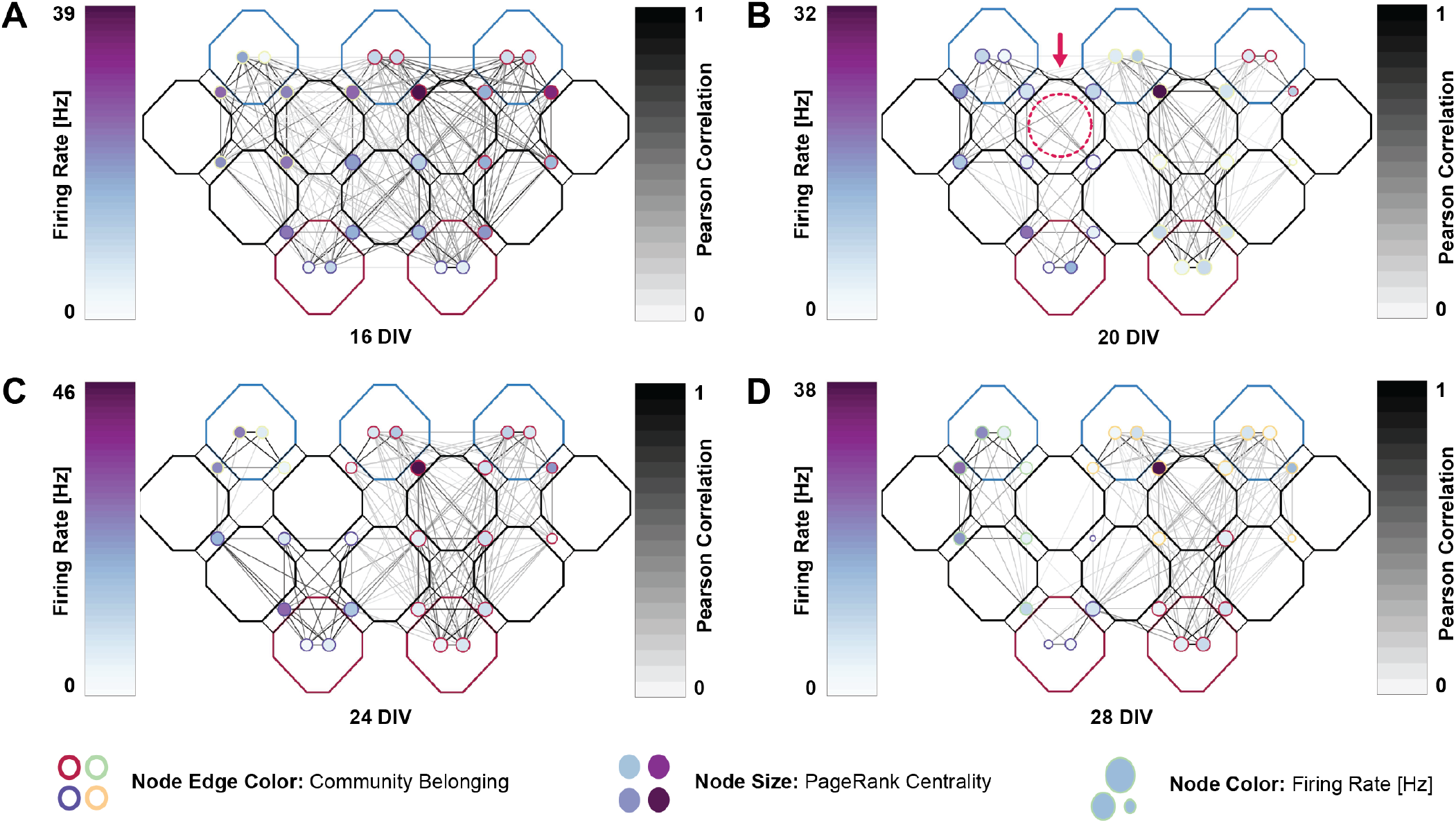
Dynamic changes in the network connectome before and after perturbation. **(A.)**-**(D.)** Representative graphs of a single 12-nodal network at 16, 20, 24 and 28 days *in vitro* (DIV). The graphs depict individual electrodes as nodes and their correlation as edges. Node color represents firing rate, node size represents PageRank centrality, and the color of edges around the nodes represents community belonging based on the Louvain algorithm. Following perturbation at 21 DIV, it is evident how functional connectivity across the hypoxic node is disrupted. Additionally, a strengthening of functional connections in uperturbed parts of the networks is observed at 24 DIV.

### The Hierarchical Microfluidic Layout Enables Investigations of Plasticity in Information Transmission Pathways Following Localized Perturbation

To evaluate plasticity changes in the networks following localized perturbation, its impact on information processing pathways going between layer 1 and 4 was evaluated (**Figure 7A**). The path length between nodes was calculated as the inverse of the correlation, summing over all paths connecting two nodes (**Figures 7B - 7G**). Path lengths increased over time, with the degree of increase contingent on the centrality of the hypoxic node in information transmission. Additionally, individual networks exhibited differential susceptibility to the perturbation, with some maintaining more robust and consistent paths between nodes in layer 1 and 4 post-perturbation. For instance, node 1 had a total of three pathways connecting it to node 10 (paths 1, 2 and 4), with two of these pathways passing directly through the targeted node (paths 2 and 4) (**Figure 7A**). The lateral pathway to the hypoxic node (path 1) maintained a consistent path length over time, both before and after perturbation (**Figure 7H**). Conversely, the paths through the hypoxic node (paths 2 and 4) exhibited increased path length over time, with an increasing number of networks showing functionally disrupted pathways as time progressed post-perturbation (**Figures 7I & 7J**). Similar results were found for the pathways connecting the other nodes (**Figure S2**). These results demonstrate the robustness of this model system for studying plasticity changes in the functional connectome following localized perturbations.

**Figure 7.**
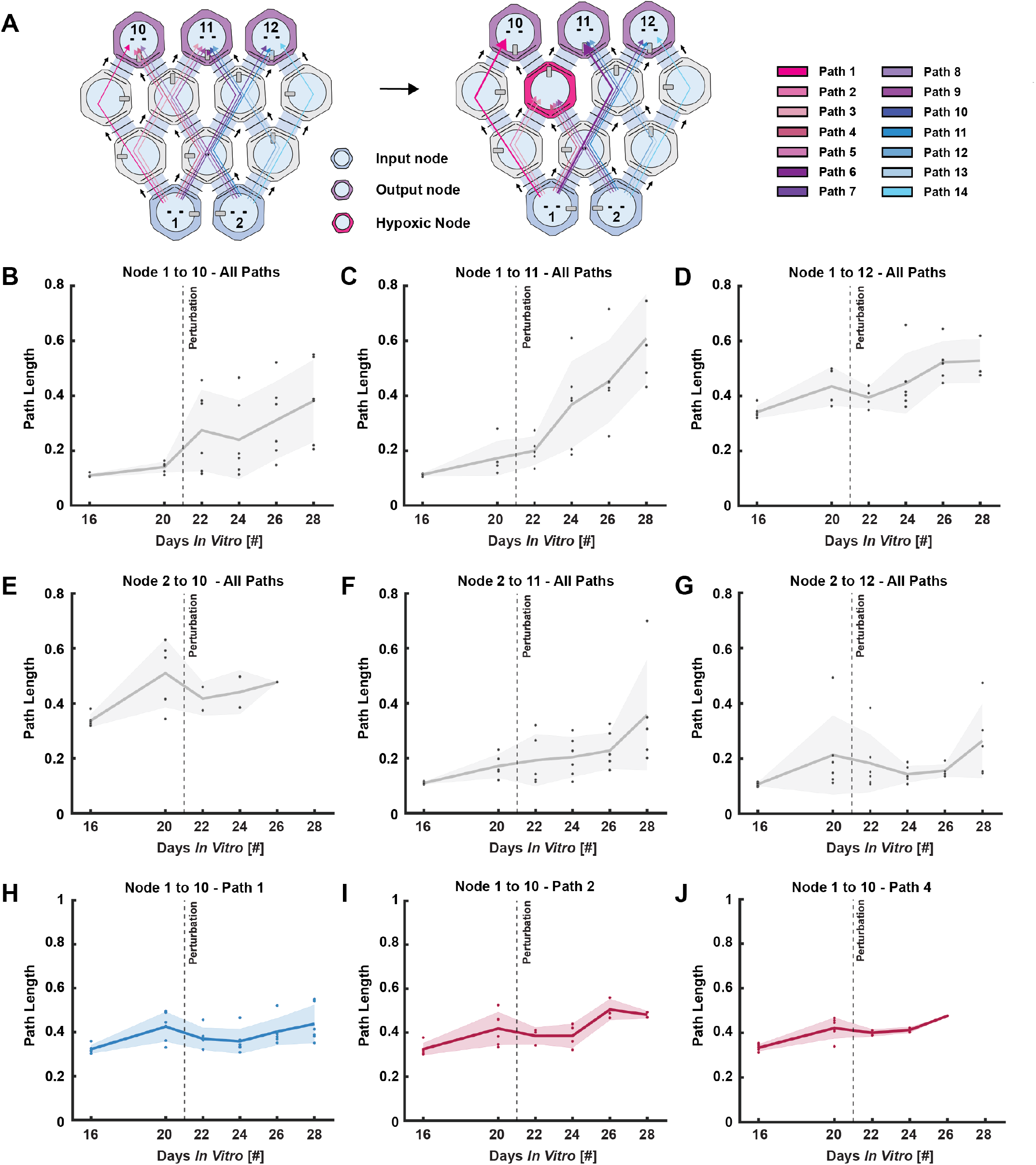
Alterations in information processing pathways following localized perturbation. **(A.)** Schematic illustrating changes in information processing pathways following perturbation in a central node of layer 3 in the 12-nodal networks. **(B.)** - **(G.)** Graphs showing alterations in path length between nodes in layer 1 and 4 of the 12-nodal cortical networks before and after localized perturbation. The impact of the perturbations varies distinctly based on the centrality of the hypoxic node for information propagation between the two interconnected nodes. **(H.)** Graph depicting the path length of the only pathway not passing through the hypoxic node connecting node 1 and 10. A slight decline in path length is observed the day after perturbation, followed by a gradual increase over time. **(I.)** - **(J.)** Graphs displaying the path length of the other two pathways connecting node 1 and node 10, going directly through the hypoxic node. A decreasing number of networks maintain functional connections through the hypoxic node over time, as reflected by the increased path length and diminishing number of data points.

### Perturbation Alters Stimulation Responsiveness Across Pathways

To assess the effects of localized perturbation on signal transmission within the networks, changes in the forward propagation of stimulation-induced activity from layer 1 to layer 4 were analysed between 20 and 28 DIV. The majority of pathways not passing through the hypoxic node maintained their sensitivity to stimulations, displaying a consistent response in the days following the localized perturbation (**Figure 8A**). In contrast, pathways that passed directly through the perturbed node showed a gradual decline in responsiveness over time (**Figures 8B & 8C**). Prior to perturbations, stimulations induced activity that propagated across all three layers of microtunnels in more than 40 % of all available pathways. By 22 DIV, one day after the perturbation, there was a 10 % increase in activated pathways going around the hypoxic node, reaching 56.2 %, while less than 3 % of pathways passing through the hypoxic node exhibited activity propagating through all three layers of microtunnels (**Figure 8D**). Although pathways going around the hypoxic nodes also showed a moderate decline in responsiveness to stimulations past 22 DIV, the extent of this decline was less pronounced compared to the pathways passing through the hypoxic node. External stimulations were also applied to all nodes in level 4 between 20 to 28 DIV, confirming maintenance of a feedforward microcircuit configuration by the absence of activity propagating beyond the first layer of microtunnels (results not shown). This analysis reveals that localized perturbations disrupted signal transmission within the networks, with pathways directly passing through the perturbed node showing a gradual decline in responsiveness over time, while pathways going around the targeted node maintained a more stable response to stimulations.

**Figure 8.**
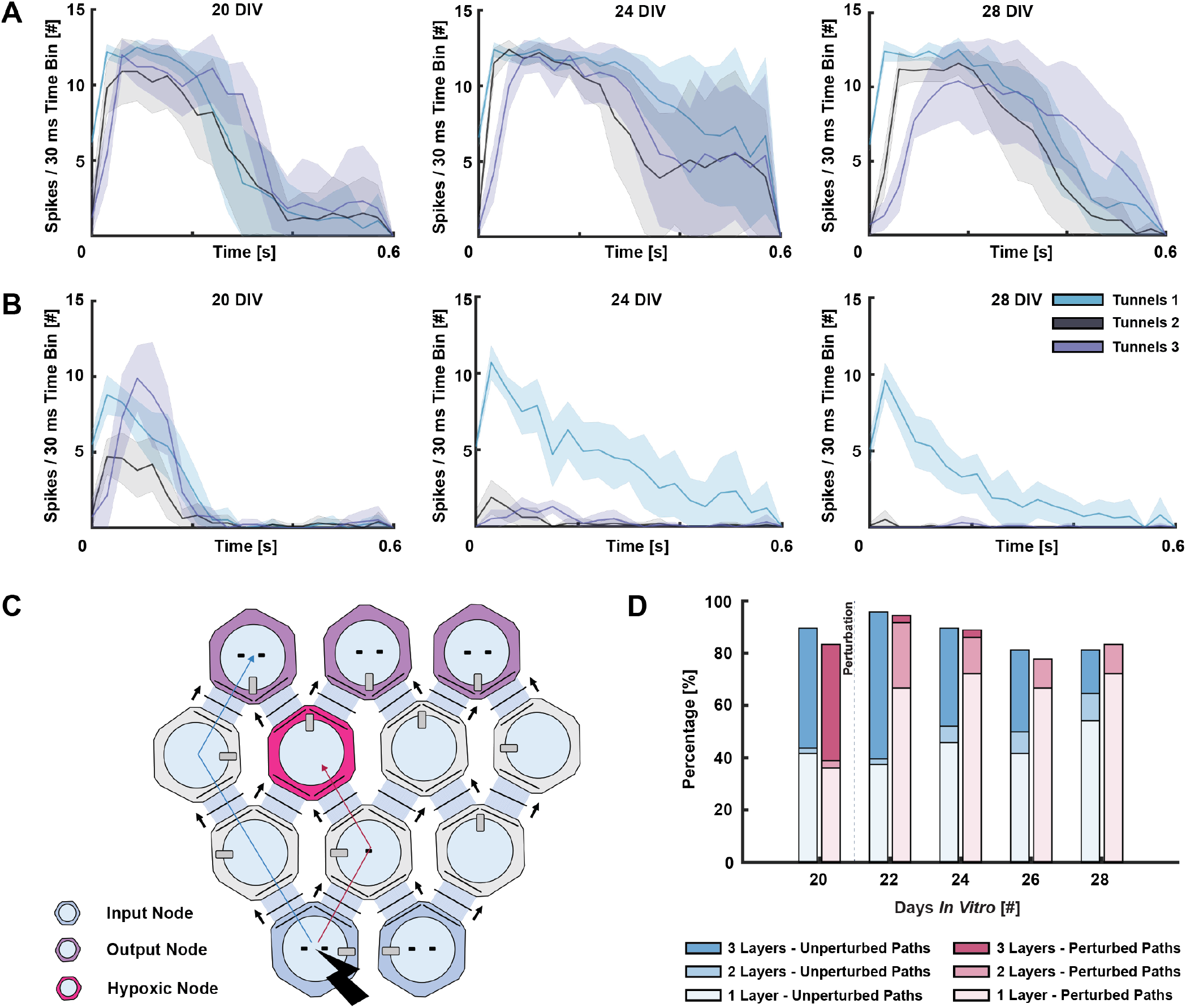
Alterations in the propagation of stimulation-evoked activity following localized perturbation. **(A.)** Peristimulus time histograms (PSTHs) displaying the average response of 10 consecutive stimulations going through an unperturbed pathway between node 1 and 10 at 20, 24 and 28 days *in vitro* (DIV). The response to stimulation remained consistent over time. **(B.)** PSTHs displaying the average response of 10 consecutive stimulations across a pathway going through the hypoxic node between node 1 and 10 at 20, 24 and 28 DIV. A progressive decrease in response is observed across tunnels 2 and 3 between 20 and 28 DIV. **(C.)** Schematic illustrating the pathway going around the hypoxic node, and the pathway passing directly through the hypoxic node in figures A and B. **(D.)** Histogram showing the percentage of stimulation-induced activity spanning 1, 2, or 3 layers of microtunnels when stimulating nodes in layer 1 over time. Blue boxes represent pathways that do not pass through the perturbed node, while pink boxes represent pathways that pass directly through the perturbed node. The propagation of stimulation-evoked activity is markedly reduced in the pathways going through the hypoxic node, in contrast to the more moderate reduction seen in pathways going around the hypoxic node.

## Discussion

In this study, we have demonstrated an advanced model system for structuring and studying hierarchical multi-nodal cortical neural networks with controlled connectivity *in vitro*. Neocortical microcircuits have a well-defined, layered structure with controlled, feedforward axonal projections connecting the distinct layers. The model system presented here recapitulates the fundamental attributes of this structure through the implementation of controlled topological constraints. We show that selective perturbation of an identified hub node within the network abolished information flow across the targeted node and significantly influenced neighboring pathways for information transmission in the multilayer networks. These results illustrate the powerful potential of neuroengineering approaches to model physiologically relevant neural microcircuits with predefined hierarchies. Such engineered neocortical circuits are highly relevant for disease modelling. A range of neurological and psychiatric disorders, among others Alzheimer’s disease, schizophrenia, stroke and Huntington’s disease, are linked to abnormalities in the neocortex (66–69). Numerous studies have shown that brain networks from patients with these conditions exhibit clear deviations from those of healthy subjects (70, 71). One such deviation is the increased vulnerability of hub areas to pathological perturbations (72–74). Over time, pathological damage to these hubs leads to overload and node failure as the node cannot handle the massive amount of incoming information, likely due to the high energetic demand required to maintain them (75). Due to their central role in network integration, failure of these hubs can thus have widespread effects on network function (25). Microfluidic models harbor substantial, albeit underutilized, potential for elucidating pathological mechanisms and propagation of relevant dynamics within controlled microenvironments (76, 77). For example, several studies have illustrated how the dysfunction of one node affects the function of connected healthy nodes in various diseases, including how misfolded protein aggregates can spread in such networks (11, 78–81). A key advantage of the presented model system is that it enables the application of localized perturbations to individual nodes within a complex network hierarchy, while allowing precise monitoring of spatiotemporal changes in neighboring areas, as demonstrated. Various types of perturbations can be applied to study their impact on network functionality, ranging from electrical (82), chemical (83), chemogenetic (84), to pharmacological (85–87). Additionally, by selectively perturbing distinct nodes within the 12-nodal interface, it is also possible to analyze how the centrality of different nodes influences network functionality and adaptation to damage. The inherent plasticity of cortical networks renders them particularly suitable for such investigations (88, 89).

Several models have been proposed to explain the changes in brain network dynamics over time in the presence of disease or trauma, yet a unified understanding remains elusive (90). Furthermore, a persistent challenge lies in the significant variability of both structural and functional alterations observed across studies (73). For instance, in Alzheimer’s disease, some studies report increased path length and decreased global efficiency with time, while others have observed the opposite (71, 91–95). This underscores the complexity of brain network alterations in response to trauma and disease, and the importance of improved model systems to study dynamic changes at early stages to understand and predict their progression and impact at multiple spatiotemporal scales. Engineering neural networks with multiple layers and pathways connecting the different nodes across these layers, as shown here, can facilitate the examination of a wide spectrum of adaptive and maladaptive responses to induced or inherent pathological perturbations. This includes the study of compensatory mechanisms such as neural reserves or degeneracy, where in the presence of an evolving perturbation, other network components can compensate for the damage (25, 96, 97). The emergence of new functional connections and the strengthening of existing ones in non-perturbed pathways three days after perturbation demonstrate that the presented model system can effectively recapitulate such compensatory mechanisms *in vitro*. While most studies to date have employed a two-nodal device featuring one healthy and one diseased population, the larger number of nodes adopted in this study offers a distinct advantage. Overall, the intricate design of the presented hierarchical platform facilitates the study of a wide range of connectomic adaptations to localized perturbations.

Microfluidic MEAs can readily be used to co-culture distinct neuronal subtypes, enhancing the physiological relevance of emerging network behaviours (11, 98–101). For this study, we used cortical embryonic rat neurons and demonstrated the fundamental attributes of this model system for engineering complex networks, identifying central hubs in the system, selectively perturbing them, and investigating the effects of the perturbation across the hierarchical network. Despite the canonical structure of cortical microcircuits, there is significant heterogeneity across the cortex, with different layers containing distinct neuronal subtypes that contribute to diverse connectivity patterns and functions (40). The engineered microcircuit shown in the present study is an excellent model system for studying physiological and pathophysiological changes in multi-layer, hierarchical neocortical networks *in vitro*. Future work may thus incorporate regionspecific cortical cells into the model, for example, for specific investigations of cortical microcircuit formation and encoding.

## Conclusion

Engineered neural networks represent a versatile tool for investigating the dynamics of neural circuits in controlled microenvironments. This study has introduced a novel microfluidic platform featuring 12 hierarchically interconnected nodes, designed to mimic the laminar organization of neocortical neural networks *in vitro*. A significant advantage with the large number of interconnected nodes is the facilitation of precise, targeted perturbations, enabling the study of adaptive compensatory mechanisms such as neural reserve and degeneracy. By inducing hypoxia in a central hub node, we have demonstrated the utility of this platform for probing plasticity changes, revealing adaptations in both directly impacted and surroundings network regions. This makes the present model highly relevant for advanced disease modelling.

## AUTHOR CONTRIBUTIONS

The author contributions follow the CRediT system. **NWH**: Conceptualization, Methodology, Software, Investigation (chip design & manufacturing, cell experiments, ICC, electrophysiology, formal analysis), Writing – Original Draft, Visualization. **PS, AS, IS**: Conceptualization, Methodology, Writing – Review & Editing, Resources, Funding Acquisition.

## ACKNOWLEDGEMENTS

We would like to thank Dr. Rajeevkumar Nair Raveendran at the Viral Vector Core Facility, Kavli Institute for systems neuroscience, for designing and preparing the AAV viruses used for structural analysis and calcium imaging. Prof. Michela Chiappalone and Prof. Sergio Martinoia, University of Genova are acknowledged for generously providing the scripts for the Precise Timing Spike Detection algorithm. Polina Malahov is acknowledged for optimizing the tools used for calcium imaging.

## Bibliography

1. L. Luo. Architectures of neuronal circuits. Science, 373(6559):eabg7285, 9 2021. doi: 10.1126/science.abg7285.

2. V. Pasquale, P. Massobrio, L. L. Bologna, M. Chiappalone, and S. Martinoia. Self-organization and neuronal avalanches in networks of dissociated cortical neurons. Neuroscience, 153(4):1354–1369, 6 2008. doi: 10.1016/j.neuroscience.2008.03.050.

3. D. A. Wagenaar, J. Pine, and S. M. Potter. An extremely rich repertoire of bursting patterns during the development of cortical cultures. BMC Neurosci., 7:11, 2 2006. doi: 10.1186/1471-2202-7-11.

4. L. N. Borodinsky, Y. H. Belgacem, I. Swapna, O. Visina, O. A. Balashova, E. B. Sequerra, M. K. Tu, J. B. Levin, K. A. Spencer, P. A. Castro, A. M. Hamilton, and S. Shim. Spatiotemporal integration of developmental cues in neural development. Dev. Neurobiol., 75 (4):349–359, 4 2015. doi: 10.1002/dneu.22254.

5. G. S. Withers, C. D. James, C. E. Kingman, H. G. Craighead, and G. A. Banker. Effects of substrate geometry on growth cone behavior and axon branching. J. Neurobiol., 66(11): 1183–1194, 9 2006. doi: 10.1002/neu.20298.

6. E. W. Dent, S. L. Gupton, and F. B. Gertler. The Growth Cone Cytoskeleton in Axon Outgrowth and Guidance. Cold Spring Harb. Perspect. Biol., 3(3):1–39, 3 2011. doi: 10.1101/CSHPERSPECT.A001800.

7. G. Gangatharan, S. Schneider-Maunoury, and M. A. Breau. Role of mechanical cues in shaping neuronal morphology and connectivity. Biol. Cell, 110(6):125–136, 6 2018. doi: 10.1111/boc.201800003.

8. A. M. Taylor, S. W. Rhee, C. H. Tu, D. H. Cribbs, C. W. Cotman, and N. L. Jeon. Microfluidic Multicompartment Device for Neuroscience Research. Langmuir, 19(5):1551–1556, 3 2003. doi: 10.1021/la026417v.

9. H. Yamamoto, S. Moriya, K. Ide, T. Hayakawa, H. Akima, S. Sato, S. Kubota, T. Tanii, M. Niwano, S. Teller, J. Soriano, and A. Hirano-Iwata. Impact of modular organization on dynamical richness in cortical networks. Sci. Adv., 4(11):eaau4914, 11 2018. doi: 10.1126/sciadv.aau491.

10. S. B. Lassers, Y. S. Vakilna, W. C. Tang, and G. J. Brewer. The flow of axonal information among hippocampal sub-regions 2: patterned stimulation sharpens routing of information transmission. Front. Neural Circuits, 17:1272925, 12 2023. doi: 10.3389/fncir.2023.1272925.

11. K. S. Hanssen, N. Winter-Hjelm, S. N. Niethammer, A. Kobro-Flatmoen, M. P. Witter, A. Sandvig, and I. Sandvig. Reverse Engineering of Feedforward Cortical-Hippocampal Microcircuits for Modelling Neural Network Function and Dysfunction. bioRxiv, 2 2024. doi: 10.1101/2023.06.26.546556.

12. M. U. Park, Y. Bae, K.-S. Lee, J. H. Song, S.-M. Lee, and K.-H. Yoo. Collective dynamics of neuronal activities in various modular networks. Lab Chip, 21(5):951–961, 3 2021. doi: 10.1039/D0LC01106A.

13. N. Winter-Hjelm, Å. B. Tomren, P. Sikorski, A. Sandvig, and I. Sandvig. Structure-function dynamics of engineered, modular neuronal networks with controllable afferent-efferent connectivity. J. Neural Eng., 20:046024, 8 2023. doi: 10.1088/1741-2552/ace37f.

14. J.-M. Peyrin, B. Deleglise, L. Saias, M. Vignes, P. Gougis, S. Magnifico, S. Betuing, M. Pietri, J. Caboche, P. Vanhoutte, J.-L. Viovy, and B. Brugg. Axon diodes for the reconstruction of oriented neuronal networks in microfluidic chambers. Lab Chip, 11(21): 3663–3673, 11 2011. doi: 10.1039/C1LC20014C.

15. A. Gladkov, Y. Pigareva, D. Kutyina, V. Kolpakov, A. Bukatin, I. Mukhina, V. Kazantsev, and A. Pimashkin. Design of Cultured Neuron Networks in vitro with Predefined Connectivity Using Asymmetric Microfluidic Channels. Sci. Rep., 7:15625, 12 2017. doi: 10.1038/s41598-017-15506-2.

16. P. M. Holloway, G. I. Hallinan, M. Hegde, S. I. R. Lane, K. Deinhardt, and J. West. Asymmetric confinement for defining outgrowth directionality. Lab Chip, 19(8):1484–1489, 4 2019. doi: 10.1039/C9LC00078J.

17. R. Renault, J.-B. Durand, J.-L. Viovy, and C. Villard. Asymmetric axonal edge guidance: a new paradigm for building oriented neuronal networks. Lab Chip, 16(12):2188–2191, 6 2016. doi: 10.1039/C6LC00479B.

18. P. Sterling and S. Laughlin. Principles of neural design. The MIT Press, 2015. ISBN 9780262534680.

19. O. Sporns. From simple graphs to the connectome: Networks in neuroimaging. NeuroImage, 62(2):881–886, 8 2012. doi: 10.1016/j.neuroimage.2011.08.085.

20. D. J. Watts and S. H. Strogatz. Collective dynamics of ‘small-world’ networks. Nature, 393: 440–442, 6 1998. doi: 10.1038/30918.

21. E. Bullmore and O. Sporns. Complex brain networks: graph theoretical analysis of structural and functional systems. Nat. Rev. Neurosci., 10:186–198, 2 2009. doi: 10.1038/nrn2575.

22. C. J. Stam and E. C. W. van Straaten. The organization of physiological brain networks. Clin Neurophysiol., 123(6):1067–1087, 6 2012. doi: 10.1016/J.CLINPH.2012.01.011.

23. S. Oldham and A. Fornito. The development of brain network hubs. Dev. Cogn. Neurosci., 36:100607, 4 2019. doi: 10.1016/j.dcn.2018.12.005.

24. E. Bullmore and O. Sporns. The economy of brain network organization. Nat. Rev. Neurosci., 13:336–349, 4 2012. doi: 10.1038/nrn3214.

25. A. Fornito, A. Zalesky, D. S. Bassett, D. Meunier, I. Ellison-Wright, M. Yücel, S. J. Wood, K. Shaw, J. O’Connor, D. Nertney, B. J. Mowry, C. Pantelis, and E. T. Bullmore. Genetic influences on cost-efficient organization of human cortical functional networks. J. Neurosci., 31(9):3261–3270, 3 2011. doi: 10.1523/jneurosci.4858-10.2011.

26. M. P. van den Heuvel and O. Sporns. Network hubs in the human brain. Trends Cogn. Sci., 17(12):683–696, 12 2013. doi: 10.1016/j.tics.2013.09.012.

27. F. V. Farahani, W. Karwowski, and N. R. Lighthall. Application of graph theory for identifying connectivity patterns in human brain networks: A systematic review. Front. Neurosci., 13: 585, 6 2019. doi: 10.3389/fnins.2019.00585.

28. P. Massobrio, V. Pasquale, and S. Martinoia. Self-organized criticality in cortical assemblies occurs in concurrent scale-free and small-world networks. Sci. Rep., 5:10578, 6 2015. doi: 10.1038/srep10578.

29. V. Fiskum, N. Winter-Hjelm, N. Christiansen, A. Sandvig, and I. Sandvig. ALS patient-derived motor neuron networks exhibit microscale dysfunction and mesoscale compensation rendering them highly vulnerable to perturbation. bioRxiv, 1 2024. doi: 10.1101/2024.01.04.574167.

30. K. Heiney, O. H. Ramstad, V. Fiskum, A. Sandvig, I. Sandvig, and S. Nichele. Neuronal avalanche dynamics and functional connectivity elucidate information propagation in vitro. Front. Neural Circuits, 16:980631, 9 2022. doi: 10.3389/fncir.2022.980631.

31. P. Rakic. Evolution of the neocortex: a perspective from developmental biology. Nat. Rev. Neurosci., 10:724–735, 10 2009. doi: 10.1038/nrn2719.

32. P. S. Goldman-Rakic. Cellular basis of working memory. Neuron, 14(3):477–485, 1995. doi: 10.1016/0896-6273(95)90304-6.

33. A. M. Bastos, W. M. Usrey, R. A. Adams, G. R. Mangun, P. Fries, and K. J. Friston. Canonical microcircuits for predictive coding. Neuron, 76(4):695–711, 11 2012. doi: 10.1016/j.neuron.2012.10.038.

34. C. Koch, M. Massimini, M. Boly, and G. Tononi. Neural correlates of consciousness: progress and problems. Nat. Rev. Neurosci., 17:307–321, 4 2016. doi: 10.1038/nrn.2016.22.

35. E. K. Miller. The prefontral cortex and cognitive control. Nat. Rev. Neurosci., 1:59–65, 2000. doi: 10.1038/35036228.

36. A. E. Papale and B. M. Hooks. Circuit changes in motor cortex during motor skill learning. Neuroscience, 368:283–297, 1 2018. doi: 10.1016/j.neuroscience.2017.09.010.

37. R. Lorente De Nó. Analysis of the activity of the chains of internunical neurons. J. Neurophysiol., 1(3):207–244, 5 1938. doi: 10.1152/jn.1938.1.3.207.

38. V. B. Mountcastle. Modality and topographic properties of single neurons of cat’s somatic sensory cortex. J. Neurophysiol., 20(4):408–434, 7 1957. doi: 10.1152/jn.1957.20.4.408.

39. D. P. Buxhoeveden and M. F. Casanova. The minicolumn hypothesis in neuroscience. Brain, 125(5):935–951, 5 2002. doi: 10.1093/brain/awf110.

40. V. B. Mountcastle. The columnar organization of the neocortex. Brain, 120(4):701–722, 1997. doi: 10.1093/brain/120.4.701.

41. I. Thompson. Cortical development: A role for spontaneous activity? Curr. Biol., 7(5): R324–R326, 5 1997. doi: 10.1016/S0960-9822(06)00150-3.

42. A. H. Leighton and C. Lohmann. The Wiring of Developing Sensory Circuits-From Patterned Spontaneous Activity to Synaptic Plasticity Mechanisms. Front. Neural Circuits, 10: 71, 9 2016. doi: 10.3389/fncir.2016.00071.

43. D. Poli, V. P. Pastore, and P. Massobrio. Functional connectivity in in vitro neuronal assemblies. Front. Neural Circuits, 9:57, 10 2015. doi: 10.3389/fncir.2015.00057.

44. M. Chiappalone, M. Bove, A. Vato, M. Tedesco, and S. Martinoia. Dissociated cortical networks show spontaneously correlated activity patterns during in vitro development. Brain Res., 1093(1):41–53, 6 2006. doi: 10.1016/j.brainres.2006.03.049.

45. T. K. Hensch. Critical period plasticity in local cortical circuits. Nat. Rev. Neurosci., 6: 877–888, 11 2005. doi: 10.1038/nrn1787.

46. T. Wieloch and K. Nikolich. Mechanisms of neural plasticity following brain injury. Curr. Opin. Neurobiol., 16(3):258–264, 6 2006. doi: 10.1016/j.conb.2006.05.011.

47. M. Sur and C. A. Leamey. Development and plasticity of cortical areas and networks. Nat. Rev. Neurosci., 2:251–262, 2001. doi: 10.1038/35067562.

48. K. N Richter, N. H Revelo, K. J Seitz, M. S Helm, D. Sarkar, R. S. Saleeb, E. D’Este, J. Eberle, E. Wagner, C. Vogl, D. F. Lazaro, F. Richter, J. Coy-Vergara, G. Coceano, E. S. Boyden, R. R. Duncan, S. W. Hell, M. A. Lauterbach, S. E. Lehnart, T. Moser, T. F. Outeiro, P. Rehling, B. Schwappach, I. Testa, B. Zapiec, and S. O. Rizzoli. Glyoxal as an alternative fixative to formaldehyde in immunostaining and super-resolution microscopy. EMBO J., 37 (1):139–159, 1 2018. doi: 10.15252/embj.201695709.

49. W. Zou, M. Yan, W. Xu, H. Huo, L. Sun, Z. Zheng, and X. Liu. Cobalt chloride induces PC12 cells apoptosis through reactive oxygen species and accompanied by AP-1 activation. J. Neurosci. Res., 64(6):646–653, 6 2001. doi: 10.1002/jnr.1118.

50. E. A. L. Raaijmakers, N. Wanders, R. M. C. Mestrom, and R. Luttge. Analyzing Developing Brain-On-Chip Cultures with the CALIMA Calcium Imaging Tool. Micromachines, 12(4): 412, 4 2021. doi: 10.3390/mi12040412.

51. F. D. W. Radstake, E. A. L. Raaijmakers, R. Luttge, S. Zinger, and J. P. Frimat. CALIMA: The semi-automated open-source calcium imaging analyzer. Comput. Methods Programs Biomed., 179:104991, 10 2019. doi: 10.1016/j.cmpb.2019.104991.

52. B. Kraus. Spike Raster Plot, 2022.

53. J. C. Lansey. Beautiful and distinguishable line colors + colormap, 2021.

54. C. A. Brewer, G. W. Hatchard, and M. A. Harrower. ColorBrewer in Print: A Catalog of Color Schemes for Maps. Cartogr. Geogr. Inf. Sci., 30(1):5–32, 1 2003. doi: 10.1559/152304003100010929.

55. A. Maccione, M. Gandolfo, P. Massobrio, A. Novellino, S. Martinoia, and M. Chiappalone. A novel algorithm for precise identification of spikes in extracellularly recorded neuronal signals. J. Neurosci. Methods, 177(1):241–249, 2 2009. doi: 10.1016/j.jneumeth.2008.09.026.

56. V. D. Blondel, J.-L. Guillaume, R. Lambiotte, and E. Lefebvre. Fast unfolding of communities in large networks. J. Stat. Mech.: Theory Exp., 2008:P10008, 10 2008. doi: 10.1088/1742-5468/2008/10/P10008.

57. M. Rubinov and O. Sporns. Complex network measures of brain connectivity: uses and interpretations. NeuroImage, 52(3):1059–1069, 9 2010. doi: 10.1016/j.neuroimage.2009.10.003.

58. D. A. Wagenaar and S. M. Potter. Real-time multi-channel stimulus artifact suppression by local curve fitting. J. Neurosci. Methods, 120(2):113–120, 10 2002. doi: 10.1016/S0165-0270(02)00149-8.

59. R. J. Mullen, C. R. Buck, and A. M. Smith. NeuN, a neuronal specific nuclear protein in vertebratesxs. Development, 116(1):201–211, 9 1992. doi: 10.1242/dev.116.1.201.

60. W. W. Schlaepfer and J. Bruce. Simultaneous up-regulation of neurofilament proteins during the postnatal development of the rat nervous system. J. Neurosci. Res., 25(1): 39–49, 1 1990. doi: 10.1002/jnr.490250106.

61. L. F. Eng, R. S. Ghirnikar, and Y. L. Lee. Glial fibrillary acidic protein: GFAP-thirty-one years (1969-2000). Neurochem. Res., 25(9-10):1439–1451, 2000. doi: 10.1023/a:1007677003387.

62. B. Wiedenmann and W. W. Franke. Identification and localization of synaptophysin, an integral membrane glycoprotein of Mr 38,000 characteristic of presynaptic vesicles. Cell, 41(3):1017–1028, 7 1985. doi: 10.1016/S0092-8674(85)80082-9.

63. K.-O. Cho, C. A. Hunt, and M. B. Kennedy. The rat brain postsynaptic density fraction contains a homolog of the drosophila discs-large tumor suppressor protein. Neuron, 9(5): 929–942, 11 1992. doi: 10.1016/0896-6273(92)90245-9.

64. J. Burtscher, R. T. Mallet, M. Burtscher, and G. P. Millet. Hypoxia and brain aging: Neurodegeneration or neuroprotection? Ageing Res. Rev., 68:101343, 7 2021. doi: 10.1016/j.arr.2021.101343.

65. V. Fiskum, A. Sandvig, and I. Sandvig. Silencing of Activity During Hypoxia Improves Functional Outcomes in Motor Neuron Networks in vitro. Front. Integr. Neurosci., 15:792863, 12 2021. doi: 10.3389/fnint.2021.792863.

66. H. Braak and E. Braak. Neuropathological stageing of Alzheimer-related changes. Acta Neuropathol., 82(4):239–259, 9 1991. doi: 10.1007/BF00308809.

67. L. D. Selemon and P. S. Goldman-Rakic. The reduced neuropil hypothesis: a circuit based model of schizophrenia. Biol. Psychiatry, 45(1):17–25, 1 1999. doi: 10.1016/S0006-3223(98)00281-9.

68. J. S. Paulsen, V. A. Magnotta, A. E. Mikos, H. L. Paulson, E. Penziner, N. C. Andreasen, and P. C. Nopoulos. Brain structure in preclinical Huntington’s disease. Biol. Psychiatry, 59(1):57–63, 1 2006. doi: 10.1016/j.biopsych.2005.06.003.

69. R. J. Nudo. Postinfarct cortical plasticity and behavioral recovery. Stroke, 38(2):840–845, 2 2007. doi: 10.1161/01.str.0000247943.12887.d2.

70. D. J. Sharp, G. Scott, and R. Leech. Network dysfunction after traumatic brain injury. Nat. Rev. Neurol., 10:156–166, 2014. doi: 10.1038/nrneurol.2014.15.

71. Y. He, Z. Chen, and A. Evans. Structural Insights into Aberrant Topological Patterns of Large-Scale Cortical Networks in Alzheimer’s Disease. J. Neurosci., 28(18):4756–4766, 4 2008. doi: 10.1523/jneurosci.0141-08.2008.

72. N. A. Crossley, A. Mechelli, J. Scott, F. Carletti, Peter T. Fox, P. Mcguire, and E. T. Bullmore. The hubs of the human connectome are generally implicated in the anatomy of brain disorders. Brain, 137(8):2382–2395, 2014. doi: 10.1093/brain/awu132.

73. B. M. Tijms, A. M. Wink, W. de Haan, W. M. van der Flier, C. J. Stam, P. Scheltens, and F. Barkhof. Alzheimer’s disease: connecting findings from graph theoretical studies of brain networks. Neurobiol. Aging, 34(8):2023–2036, 8 2013. doi: 10.1016/j.neurobiolaging.2013.02.020.

74. M. P. van den Heuvel, I. L. C. van Soelen, C. J. Stam, R. S. Kahn, D. I. Boomsma, and H. E. H. Pol. Genetic control of functional brain network efficiency in children. Eur. Neuropsychopharmacol., 23(1):19–23, 1 2013. doi: 10.1016/j.euroneuro.2012.06.007.

75. C. J. Stam. Modern network science of neurological disorders. Nat. Rev. Neurosci., 15: 683–695, 9 2014. doi: 10.1038/nrn3801.

76. J. A. del Rio and I. Ferrer. Potential of Microfluidics and Lab-on-Chip Platforms to Improve Understanding of “prion-like” Protein Assembly and Behavior. Front. Bioeng. Biotechnol., 8:570692, 9 2020. doi: 10.3389/fbioe.2020.570692.

77. T. Osaki, Y. Shin, V. Sivathanu, M. Campisi, and R. D. Kamm. In Vitro Microfluidic Models for Neurodegenerative Disorders. Adv. Healthc. Mater., 7(2):1700489, 1 2018. doi: 10.1002/adhm.201700489.

78. A. Virlogeux, E. Moutaux, W. Christaller, A. Genoux, J. Bruyère, E. Fino, B. Charlot, M. Cazorla, and F. Saudou. Reconstituting Corticostriatal Network on-a-Chip Reveals the Contribution of the Presynaptic Compartment to Huntington’s Disease. Cell Rep., 22(1):110–122, 1 2018. doi: 10.1016/J.CELREP.2017.12.013.

79. V. D. Valderhaug, O. H. Ramstad, R. van de Wijdeven, K. Heiney, S. Nichele, A. Sandvig, and I. Sandvig. Micro-and mesoscale aspects of neurodegeneration in engineered human neural networks carrying the LRRK2 G2019S mutation. Front. Cell. Neurosci., 18: 1366098, 4 2024. doi: 10.3389/fncel.2024.1366098.

80. E. C. Freundt, N. Maynard, E. K. Clancy, S. Roy, L. Bousset, Y. Sourigues, M. Covert, R. Melki, K. Kirkegaard, and M. Brahic. Neuron-to-neuron transmission of α-synuclein fibrils through axonal transport. Ann. Neurol., 72(4):517–524, 10 2012. doi: 10.1002/ana.23747.

81. G. I. Hallinan, M. Vargas-Caballero, J. West, and K. Deinhardt. Tau Misfolding Efficiently Propagates between Individual Intact Hippocampal Neurons. J. Neurosci., 39(48):9623–9632, 2019. doi: 10.1523/jneurosci.1590-19.2019.

82. D. A. Wagenaar, R. Madhavan, J. Pine, and S. M. Potter. Controlling bursting in cortical cultures with closed-loop multi-electrode stimulation. J. Neurosci., 25(3):680–688, 1 2005. doi: 10.1523/jneurosci.4209-04.2005.

83. P. Massobrio, C. N. G. Giachello, M. Ghirardi, and S. Martinoia. Selective modulation of chemical and electrical synapses of Helix neuronal networks during in vitro development. BMC Neurosci., 14:22, 2 2013. doi: 10.1186/1471-2202-14-22.

84. J. S. Weir, N. Christiansen, A. Sandvig, and I. Sandvig. Selective inhibition of excitatory synaptic transmission alters the emergent bursting dynamics of in vitro neural networks. Front. Neural Circuits, 17:1020487, 2023. doi: 10.3389/fncir.2023.1020487.

85. M. Frega, V. Pasquale, M. Tedesco, M. Marcoli, A. Contestabile, M. Nanni, L. Bonzano, G. Maura, and M. Chiappalone. Cortical cultures coupled to Micro-Electrode Arrays: A novel approach to perform in vitro excitotoxicity testing. Neurotoxicol. Teratol., 34(1):116– 127, 1 2012. doi: 10.1016/j.ntt.2011.08.001.

86. A. Odawara, H. Katoh, N. Matsuda, and I. Suzuki. Physiological maturation and drug responses of human induced pluripotent stem cell-derived cortical neuronal networks in long-term culture. Sci. Rep., 6:26181, 5 2016. doi: 10.1038/srep26181.

87. I. Colombi, S. Mahajani, M. Frega, L. Gasparini, and M. Chiappalone. Effects of antiepileptic drugs on hippocampal neurons coupled to micro-electrode arrays. Front. Neuroeng., 6: 10, 11 2013. doi: 10.3389/fneng.2013.00010.

88. J. van Pelt, I. Vajda, P. S. Wolters, M. A. Corner, and G. J. A. Ramakers. Dynamics and plasticity in developing neuronal networks in vitro. Prog. Brain Res., 147:171–188, 2005. doi: 10.1016/S0079-6123(04)47013-7.

89. P. Massobrio, J. Tessadori, M. Chiappalone, and M. Ghirardi. In vitro studies of neuronal networks and synaptic plasticity in invertebrates and in mammals using multielectrode arrays. Neural Plast., 2015:196195, 2015. doi: 10.1155/2015/196195.

90. R. Ashish and P. Fon. Models of Network Spread and Network Degeneration in Brain Disorders. Biol. Psychiatry Cogn. Neurosci. Neuroimaging, 3(9):788–797, 9 2018. doi: 10.1016/j.bpsc.2018.07.012.

91. C.-Y. Lo, P.-N. Wang, K.-H. Chou, J. Wang, Y. He, and C.-P. Lin. Diffusion tensor tractography reveals abnormal topological organization in structural cortical networks in Alzheimer’s disease. J. Neurosci., 30(50):16876–16885, 12 2010. doi: 10.1523/jneurosci.4136-10.2010.

92. C. J. Stam, B. F. Jones, G. Nolte, M. Breakspear, and P. Scheltens. Small-world networks and functional connectivity in Alzheimer’s disease. Cereb. Cortex., 17(1):92–99, 1 2007. doi: 10.1093/CERCOR/BHJ127.

93. E. J. Sanz-Arigita, M. M. Schoonheim, J. S. Damoiseaux, S. A. R. B. Rombouts, E. Maris, F. Barkhof, P. Scheltens, and C. J. Stam. Loss of ‘Small-World’ Networks in Alzheimer’s Disease: Graph Analysis of fMRI Resting-State Functional Connectivity. PLoS One, 5(11): e13788, 2010. doi: 10.1371/journal.pone.0013788.

94. C. J. Stam, W. de Haan, A. Daffertshofer, B. F. Jones, I. Manshanden, A. M. van Cappellen van Walsum, T. Montez, J. P. A. Verbunt, J. C. de Munck, B. W. van Dijk, H. W. Berendse, and P. Scheltens. Graph theoretical analysis of magnetoencephalographic functional connectivity in Alzheimer’s disease. Brain, 132(1):213–224, 2009. doi: 10.1093/brain/awn262.

95. W. de Haan, Y. A. L. Pijnenburg, R. L. M. Strijers, Y. van der Made, W. M. van der Flier, P. Scheltens, and C. J. Stam. Functional neural network analysis in frontotemporal dementia and Alzheimer’s disease using EEG and graph theory. BMC Neurosci., 10:101, 8 2009. doi: 10.1186/1471-2202-10-101.

96. A. Avena-Koenigsberger, B. Misic, and O. Sporns. Communication dynamics in complex brain networks. Nat. Rev. Neurosci., 19:17–33, 12 2018. doi: 10.1038/nrn.2017.149.

97. U. Noppeney, K. J. Friston, and C. J. Price. Degenerate neuronal systems sustaining cognitive functions. J. Anat., 205(6):433–442, 2004. doi: 10.1111/j.0021-8782.2004.00343.x.

98. C. H. Chang, T. Furukawa, T. Asahina, K. Shimba, K. Kotani, and Y. Jimbo. Coupling of in vitro Neocortical-Hippocampal Coculture Bursts Induces Different Spike Rhythms in Individual Networks. Front. Neurosci., 16:873664, 5 2022. doi: 10.3389/fnins.2022.873664.

99. S. Han, S. Bang, H. N. Kim, N. Choi, and S. H. Kim. Modulating and monitoring the functionality of corticostriatal circuits using an electrostimulable microfluidic device. Mol. Brain, 16:13, 12 2023. doi: 10.1186/S13041-023-01007-z.

100. S. Dauth, B. M. Maoz, S. P. Sheehy, M. A. Hemphill, T. Murty, M. K. Macedonia, A. M. Greer, B. Budnik, and K. K. Parker. Neurons derived from different brain regions are inherently different in vitro: A novel multiregional brain-on-a-chip. J. Neurophysiol., 117(3): 1320–1341, 3 2017. doi: 10.1152/jn.00575.2016.

101. Y. S. Vakilna, W. C. Tang, B. C. Wheeler, and G. J. Brewer. The Flow of Axonal Information Among Hippocampal Subregions: 1. Feed-Forward and Feedback Network Spatial Dynamics Underpinning Emergent Information Processing. Front. Neural Circuits, 15: 660837, 8 2021. doi: 10.3389/fncir.2021.660837.

