## Supplementary Materials for "Engineered Cortical Microcircuits for Investigations of Neuroplasticity"

**Nicolai Winter-Hjelm[1,✉], Pawel Sikorski[2], Axel Sandvig[1,3], and Ioanna Sandvig[1,✉]**

[1]Department of Neuromedicine and Movement Science, Faculty of Medicine and Health Sciences, Norwegian University of Science and Technology (NTNU), Norway
[2]Department of Physics, Faculty of Natural Sciences, Norwegian University of Science and Technology (NTNU), Trondheim, Norway
[4]Department of Neurology and Clinical Neurophysiology, St Olav´s University Hospital, Trondheim, Norway

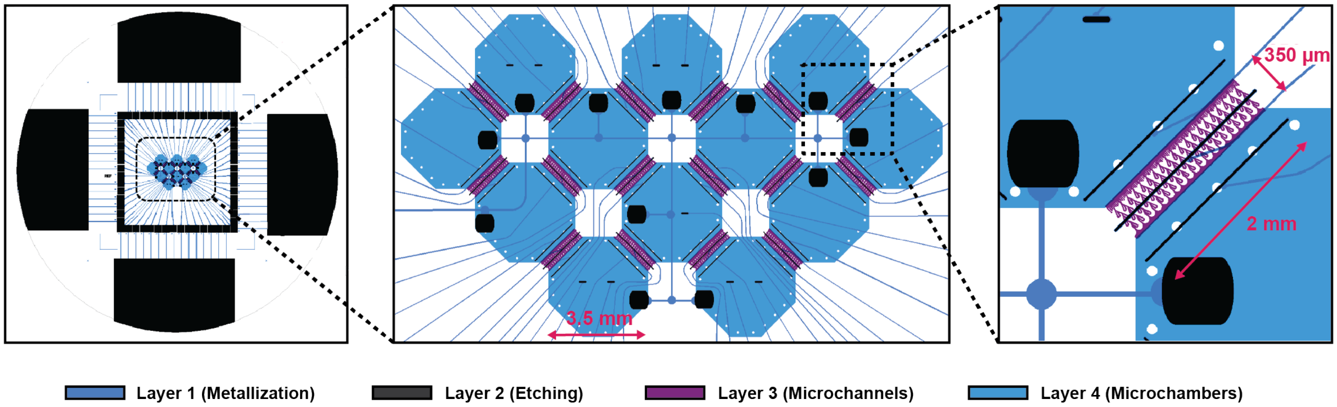


**Figure S1 | CAD design for the 12-nodal Microfluidic MEA.** Design used to fabricate the microfluidic MEAs. The four large contact pads along the edges of the wafer were used for connecting the MEAs to the potentiostat during depositions of nanoporous platinum. Following depositions, the wafers were subsequently cut into 49 x 49 mm squares with a wafer saw along the four corner marks, hence disconnecting the 60 smaller contact pads from the larger ones. Layer 3 and 4 were used for preparing the microtunnels and chambers of the microfluidic chips, respectively.

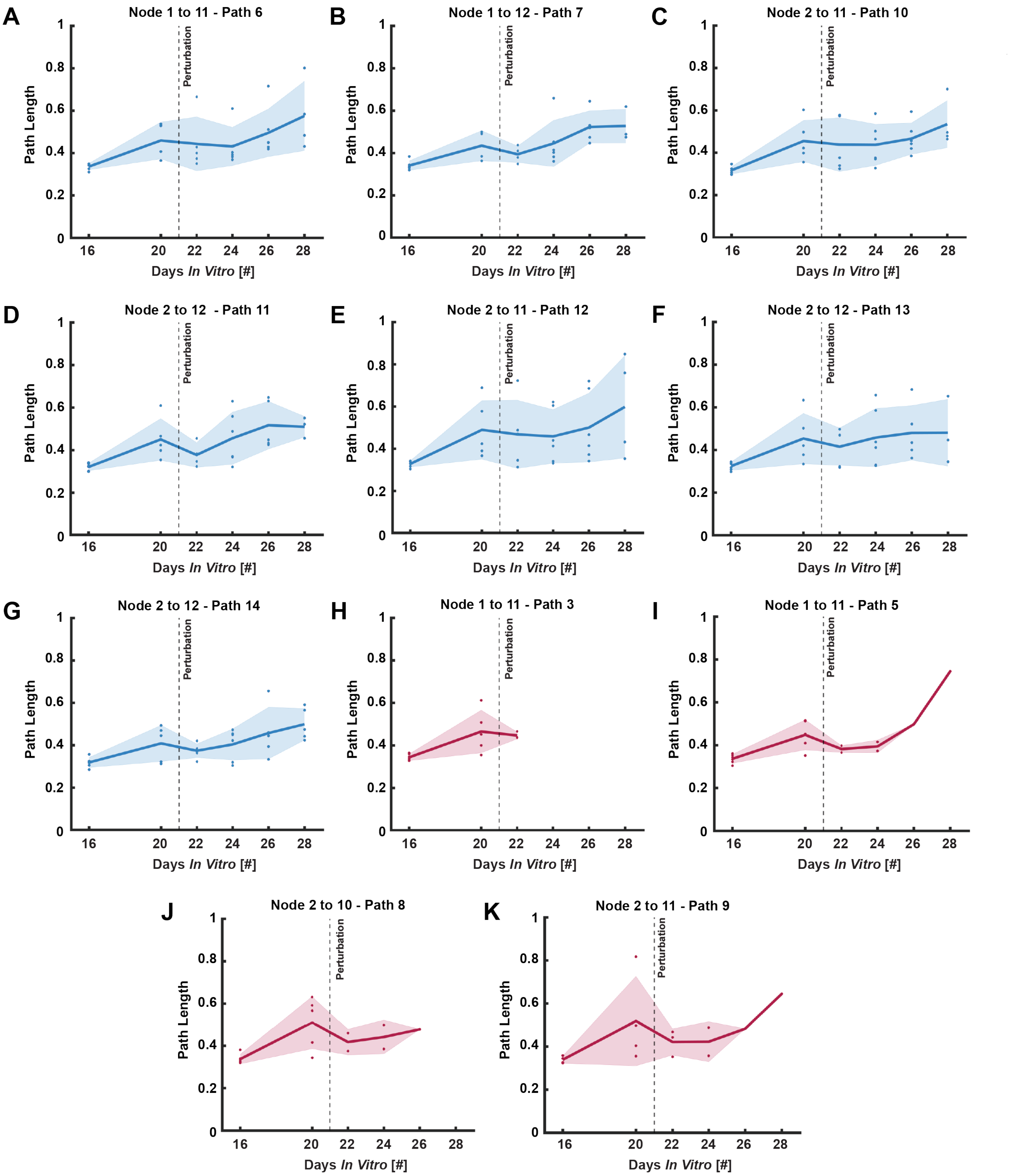


**Figure S2** | **Alterations in information processing pathways following localized induction of hypoxia**. **(A.)** - **(G.)** Graphs depicting the path length of pathways not passing directly through the hypoxic node over time. A slight decline in path length is observed the day after perturbation, followed by a gradual increase over time. **(H.)** - **(K.)** Graphs displaying the path length of pathways passing directly through the hypoxic node over time. A decreasing number of networks maintained a functional connection through the affected node over time, as reflected by the increased path length and diminishing number of data points.
